# Nanoparticle-mediated delivery of peptide-based degraders enables targeted protein degradation

**DOI:** 10.1101/2024.03.17.584721

**Authors:** Souvik Ghosal, Cara Robertus, Jeanette Wang, Harrison W. Chan, Azmain Alamgir, Joshua Almonte, Christopher A. Alabi

## Abstract

The development of small molecule-based degraders against intracellular protein targets is a rapidly growing field that is hindered by the limited availability of high-quality small molecule ligands that bind to the target of interest. Despite the feasibility of designing peptide ligands against any protein target, peptide-based degraders still face significant obstacles such as, limited serum stability and poor cellular internalization. To overcome these obstacles, we repurposed lipid nanoparticle (LNP) formulations to facilitate the delivery of Peptide-based proteolysis TArgeting Chimeras (PepTACs). Our investigations reveal robust intracellular transport of PepTAC-LNPs across various clinically relevant human cell lines. Our studies also underscore the critical nature of the linker and hydrophobic E3 binding ligand for efficient LNP packaging and transport. We demonstrate the clinical utility of this strategy by engineering PepTACs targeting two critical transcription factors, β-catenin and CREPT (cell-cycle-related and expression-elevated protein in tumor), involved in the Wnt-signalling pathway. The PepTACs induced target-specific protein degradation and led to a significant reduction in Wnt-driven gene expression and cancer cell proliferation. Mouse biodistribution studies revealed robust accumulation of PepTAC-LNPs in the spleen and liver, among other organs, and PepTACs designed against β-catenin and formulated in LNPs showed a reduction in β-catenin levels in the liver. Our findings demonstrate that LNPs can be formulated to encapsulate PepTACs, thus enabling robust delivery and potent intracellular protein degradation.

---

The past decade has seen a surge in the development of protein degrading molecules, especially those targeting various onco-proteins.^1-11^ PROTACs (PROteolysis TArgeting Chimeras) are heterobifunctional degraders that facilitate the selective degradation of a diverse array of intracellular proteins by hijacking the cell’s own naturally occurring protein degradation mechanism - the ubiquitin-proteasome system (UPS). By employing a ligand that recruits an E3 ubiquitin ligase and another that binds a protein of interest (POI), PROTACs induce an artificial interaction between an E3 and a POI. The ternary complex catalyzes selective ubiquitin-tagging of the POI, leading to its proteasome-mediated degradation. In contrast to the conventional stoichiometric binding mechanism employed by classic small molecule inhibitors, PROTACs exploit an event-driven approach that promotes the degradation of multiple POIs, ensuring a high turnover frequency and potent catalytic activity. However, a significant challenge remains,^12,13^ as many intracellular proteins are still considered “undruggable” due to the absence of a well-defined binding pocket.^14,15^

As an alternative to small molecule-based approaches, peptide-based ligands possess large protein-protein interaction surfaces, making them suitable for targeting any POI. Coupled with the rapid development of structural biology techniques that provide detailed protein-protein structural information,^16,17^ mature directed-evolution technologies such as phage and yeast display,^18-20^ and emerging computational approaches for rapid discovery of synthetic binding peptides,^6,21-23^ peptide-based ligands are ideal for extending the scope of PROTACs to “undruggable” proteins.^24,25^ Several Peptide-based Proteolysis Targeting Chimeras (PepTACs) targeting oncoproteins and transcription factors make use of a cationic cell-penetrating peptides^26,27^, cyclic peptides,^28,29^ or peptide stapling^30-33^ to facilitate cellular uptake. While promising, these approaches still suffer from poor cellular permeability, thus requiring very high doses in the tens of micromolar range for PepTAC activity. These examples are in stark contrast to small molecule PROTACs that routinely enable degradation in the sub-nanomolar range.^5,34-37^ Furthermore, PepTACs suffer from limited serum stability, preventing their wide-scale adoption in vivo.

Given the superior attributes of peptide-based ligands over small molecules, our aim was to improve the extracellular stability and cell-membrane transport of PepTACs by encapsulating them in a nanoparticle carrier (Figure 1). We envisioned a system in which nanoparticles shield the PepTAC cargo from extracellular proteases, promote cellular uptake, facilitate PepTAC escape from endosomes, and thereby enable PepTAC-mediated ubiquitination of target proteins followed by degradation through the UPS (Figure 1). We selected ionizable lipid nanoparticles (LNPs) as a carrier given the success of Patisiran,^38^ the first FDA-approved liposomal siRNA therapeutic, and recently, the COVID-19 mRNA vaccines developed by Pfizer/BioNTech and Moderna.^39,40^ We then used a previously reported 27-mer PepTAC designed against a validated onco-target, CREPT (cell-cycle-related and expression-elevated protein in tumor) to test peptide delivery capabilities of reformulated LNPs. Our studies show that PepTAC amphiphilicity is essential for LNP loading and intracellular delivery irrespective of the choice of the ionizable lipid component. PepTAC-LNP formulations against CREPT were rapidly taken up and enabled potent targeted degradation of the CREPT protein at sub-nanomolar doses across various cell lines, rivalling traditional small molecule PROTACs. To broaden the scope of this work, we designed a new PepTAC against another oncoprotein, β-catenin, and showed similar potent in vitro degradation and downstream effects on transcriptional activity. Furthermore, we assessed their potential for clinical translation and observed that systemically administered LNPs increased PepTAC accumulation in the liver and spleen, enabling β-catenin degradation. Together, our findings emphasize the influence of lipid formulation and peptide amphiphilicity on uptake efficiency and open the door for the use of PepTACs as therapeutics as well as tools for chemical biology.

**Figure 1.**
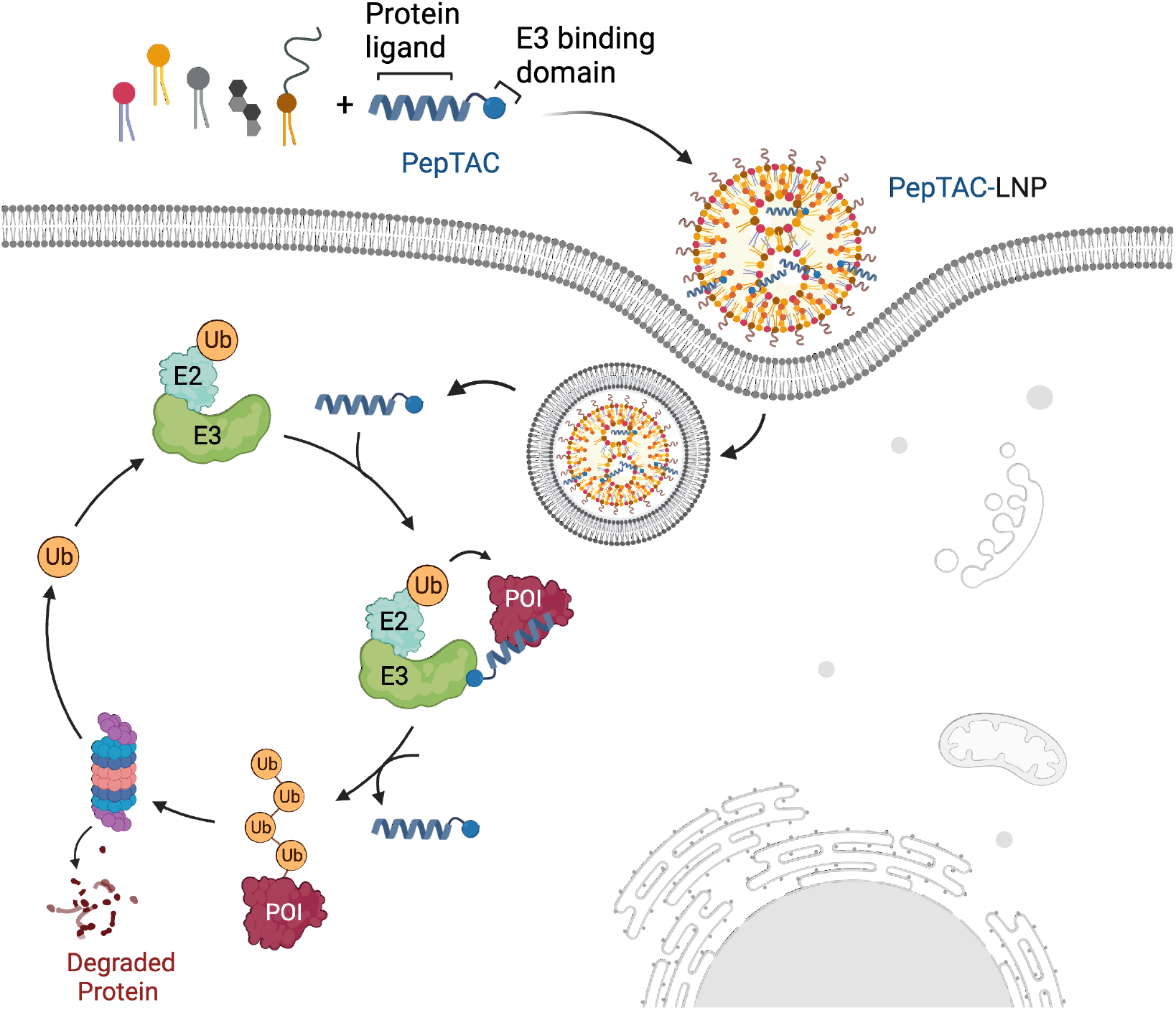
Encapsulation of PepTACs in LNPs allows for endosomal uptake, escape, and cytosolic delivery. Upon release in the cytoplasm, PepTACs form a ternary complex with the E3 ubiquitin ligase, which initiates ubiquitination and degradation of the POI by the proteosome.

## RESULTS

### Intracellular delivery of PepTACs via LNPs

We began our investigation with a PepTAC previously shown to degrade CREPT, also known as RPRD1B (regulation of nuclear pre-mRNA domain-containing protein 1B).^26^ CREPT accelerates the cell cycle and promotes tumor growth. Aberrant expression has been found in numerous types of cancers and its expression has been shown to negatively correlate with patient outcomes.^41-43^ Studies have demonstrated that CREPT interacts with RNA polymerase II, which promotes chromatin loop formation and activates cyclin D1 transcription, among others, in response to Wnt signaling.^43,44^ The PepTAC utilized in the study consisted of a 21-mer CREPT-binding sequence connected to an E3 binding peptide via a short 6-aminohexanoic acid (Ahx) linker (Figure 2A).^26^ This peptide-binding ligand was derived from a leucine-zipper-like motif located within the CREPT CCT (coiled-coil terminus) domain, specifically spanning from K266 to V286. Previous predictions using Schrödinger on CREPT CCT monomer determined that this peptide ligand binds to the same K266-V286 region, facilitated by three leucine residues.^26^ AlfaFold-Multimer predictions of the CREPT CCT dimer (PDB: 4NAD) with the peptide ligand confirmed the formation of a head-to-tail CREPT CCT homodimer and association of the peptide ligand with the leucine zipper motif at K266 to V286 (Figure 2A). In the study by Ma *et. al*., a short cell-penetrating RRRRK sequence was attached to the C-terminal of the PepTAC to enable its cellular internalization. This PepTAC, referred to as PRTC in the previous study, enabled the degradation of CREPT in PANC-1 cells at concentrations in excess of 10 µM.^26^ Using this as a starting point, we formulated the CREPT PepTAC (^CR^PepTAC) using the standard LNP formulation consisting of an ionizable MC3 lipid, cholesterol, DSPC, and DMG-PEG2K lipids at a 50:38.5:10:1.5 ratio. This formulation produced stable LNPs, as characterized by dynamic light scattering (Supplementary Figure S1). Although the addition of this PepTAC-LNP formulation to HeLa cells resulted in some cell transfection (Figure 2B), we observed that the addition of 10 mol% DOTAP as a 5^th^ lipid to this formulation resulted in robust transfection of the entire cell population (Figure 2B). This dependency on DOTAP persisted irrespective of the ionizable lipid composition. A similar trend was observed with and without DOTAP using the principal ionizable lipid in the Pfizer/BioNTech (ALC0315) and Moderna (SM102) mRNA vaccine formulations (Figure 2B). MC3 demonstrated improved ^CR^PepTAC delivery compared to ALC0315 and SM102 (Figure 2B), making it the preferred choice for subsequent transfection studies. LNP formulations with DOTAP resulted in stable nanoparticles similar to those without DOTAP, as characterized by dynamic light scattering and zeta potential (Supplementary Figures S1 and S2). The PepTAC-LNPs analyzed by cryo-electron microscopy exhibited a cup-shaped morphology (likely due to dehydration defects of the drying process) (Supplementary Figure S3). The impact of DOTAP on transfection efficiency is partially attributed to enhanced PepTAC encapsulation, exhibiting a 70% encapsulation with DOTAP versus 32% without DOTAP (Supplementary Figure S4).

**Figure 2.**
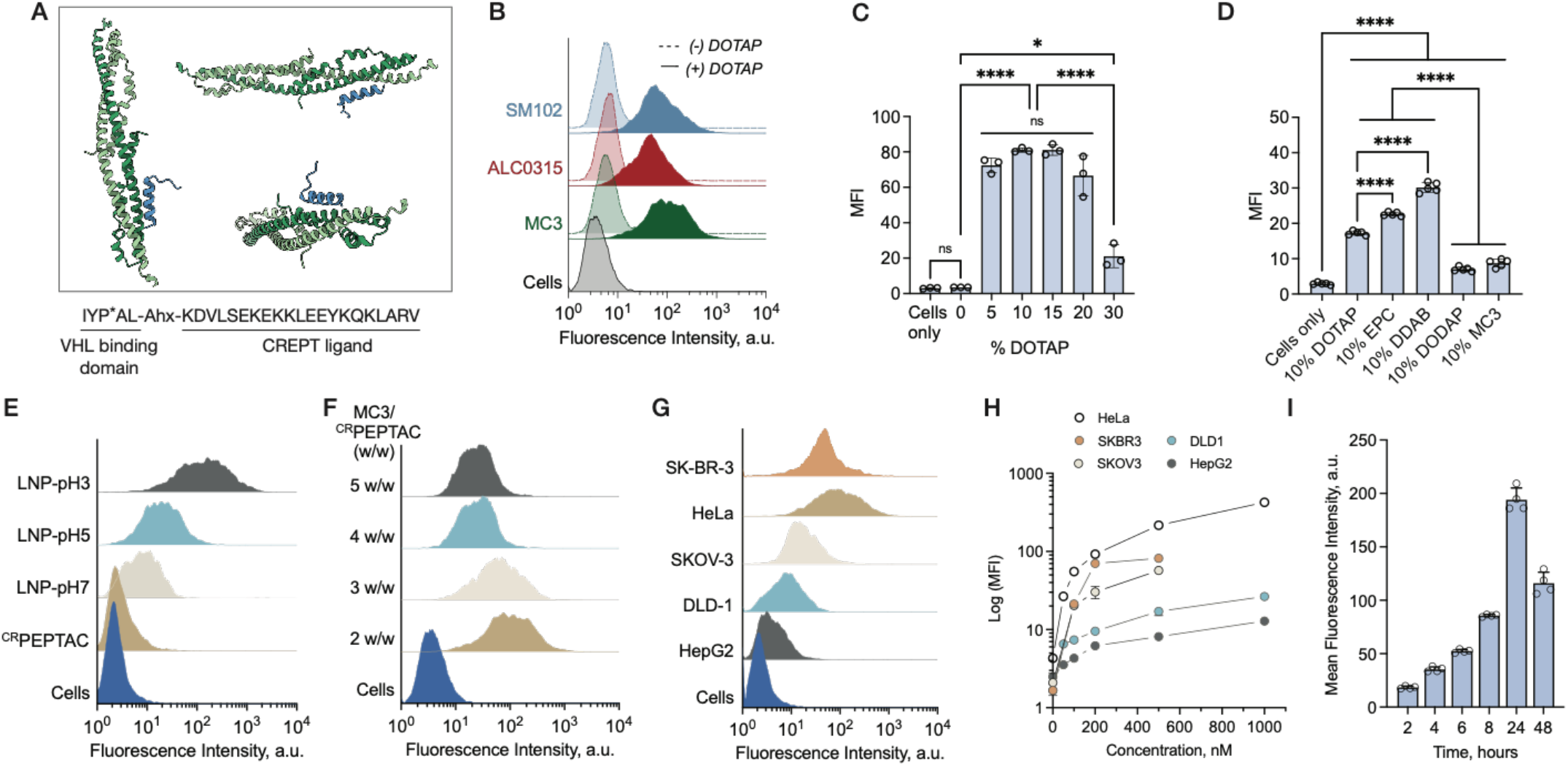
(A) AlfaFold-Multimer prediction of CREPT CCT dimer (4NAD, light and dark green) with the 21-mer CREPT ligand (blue). (B). Flow cytometry data showing transfection of LNPs formulated at pH 3 with and without DOTAP in HeLa cells at 500 nM.(C) MC3 LNPs formulated at different mol percentages of DOTAP transfected into HeLa cells with 500 nM ^CR^PepTAC. (D) Effect of different DOTAP derivatives (all at 10 mol %) on LNP-mediated ^CR^PepTAC (200 nM) delivery into HeLa cells. Effect of pH (E), and ionizable lipid/^CR^PepTAC wt/wt ratio (F) on LNP-mediated ^CR^PepTAC delivery. Delivery of ^CR^PepTAC via MC3 LNP into different cell types (G) at different concentrations (H). (I) Uptake of fluorescein-^CR^PepTAC into HeLa cells as a function of time. Data are displayed as mean ± SD by one-way ANOVA followed by Tukey correction for multiple comparisons. * p < 0.05, ** p < 0.01, ***p < 0.001, ****p < 0.0001

Formulations containing varying amounts of DOTAP exhibited minimal differences in transfection efficiency within the range of 5-20% DOTAP (Figure 2C). However, transfection efficiency declined at higher concentrations of DOTAP. The impact of DOTAP cannot be attributed to the increase in the fraction of cationic lipid relative to the other lipids in the formulation. This is evident as replacing DOTAP with an equivalent amount of MC3, or its ionizable equivalent, DODAP, led to a marked decrease in transfection efficiency relative to that seen with an equivalent amount of DOTAP (Figure 2D).

To investigate the chemical and structural characteristics of DOTAP that contribute to enhanced transfection, we examined additional structural analogues such as DDAB (Supplementary Figure S5), which possesses a different saturated lipid tail but a similar cationic headgroup, and EPC (Supplementary Figure S5), an analogue that incorporates a phosphodiester in the headgroup. The collective results show that only structural analogues with a permanently cationic headgroup improved transfection efficiency (Figure 2D). We next investigated the effect of formulation pH and lipid/ ^CR^PepTAC ratio on transfection efficiency. Transfection efficiency improved with decreasing both pH and MC3/^CR^PepTAC ratio (Figure 2E and 2F). The optimal conditions for maximum transfection efficiency were achieved with a formulation at pH3 and MC3/^CR^PepTAC ratio of 2/1. The effect of pH on transfection efficiency can be attributed in part to the ^CR^PepTAC encapsulation efficiency which showed modest encapsulation of 70% and 67% at pH 3 and 5 respectively, and poor encapsulation of only 22% at pH 7 (Supplementary Figure S4). The ^CR^PepTAC maintained its alpha-helix structure at all pHs (Figure S6). Changes to the type of phospholipid (DSPC vs. DOPE), or the phospholipid to ionizable lipid ratio, had minimal effect on cellular uptake (Supplementary Figures S7 and S8). In addition to the studies performed in HeLa cells, the optimized ^CR^PepTAC-LNP formulation also exhibits dose-dependent transfection across a diverse range of cell lines including human breast cancer (SK-BR-3), ovarian cancer (SKOV-3), colorectal adenocarcinoma (DLD-1), and hepatocellular carcinoma (HepG2) cell lines (Figure 2G and 2H, Supplementary Figures S10-S15). A similar dose-dependent uptake was also observed when the LNPs were formulated at pH 5 (Supplementary Figure S9). Cellular uptake of ^CR^PepTAC-LNP shows a rapid increase within the first 24 hours, followed by a decline in the subsequent 24 hours that may be attributed to peptide degradation and exocytosis of the free dye (Figure 2I). Cellular uptake was observed to be temperature-dependent, indicative of an energy dependent endocytic uptake mechanism (Supplementary Figure S16).

Subcellular localization of the delivered PepTACs was examined using confocal imaging (Figure 3A). ^CR^PepTAC-LNPs formulated at pH3 colocalized with an endosomal marker and showed diffuse staining indicative of cytosolic localization (Figure 3A and 3B). Conversely, the pH5 formulation displayed a higher degree of colocalization between ^CR^PepTAC and endosomes (Figure 3A). Notably, ^CR^PepTAC alone showed no uptake or cellular association. This data suggests LNPs mediate PepTAC uptake via the endosomal route, escape the endosome and subsequently release PepTACs into the cytosol.

**Figure 3.**
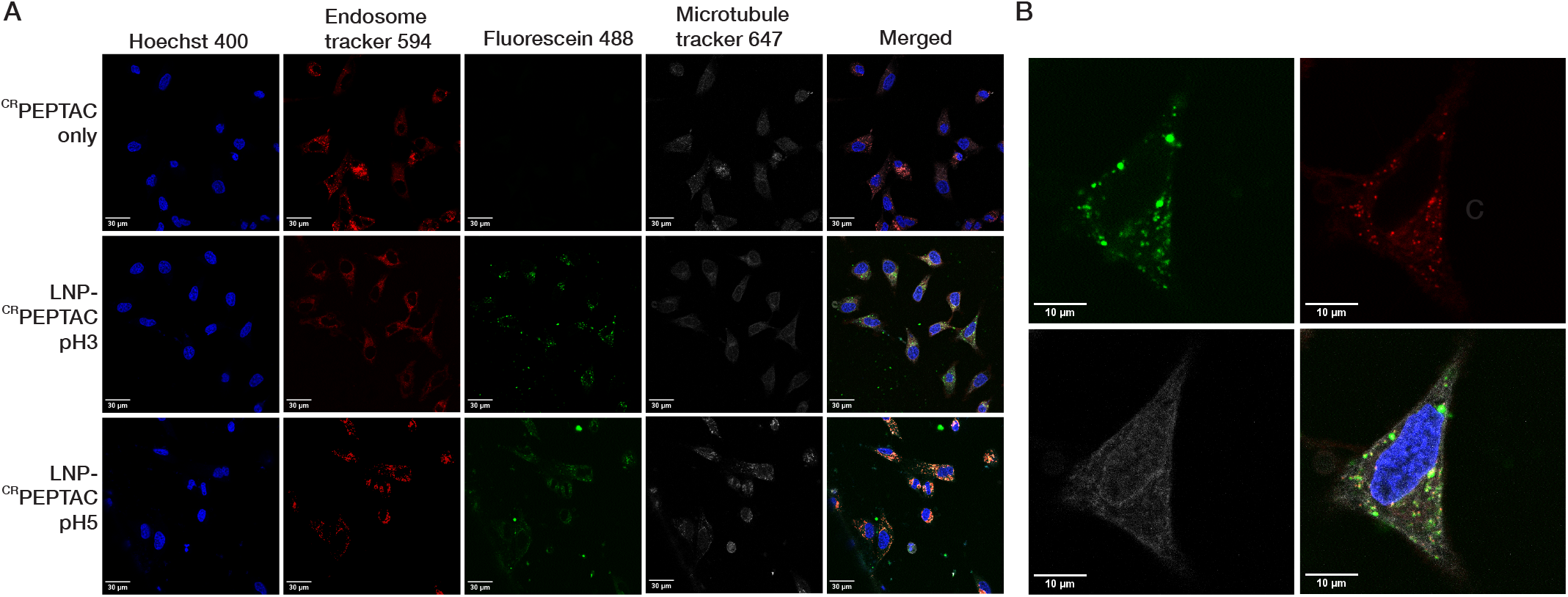
(A) Confocal imaging of LNPs with ^CR^PepTAC transfected into HeLa cells. Nucleus stained with Hoechst400 (blue), endosome tracker for endosomal staining (red), fluorescein labelled ^CR^PepTAC (green) and microtubule tracker (gray). (B) Magnified image of a single cell showing endosomal and cytoplasmic staining of fluorescein-^CR^PepTAC using LNPs formulated at pH3.

### PepTAC amphiphilicity enables encapsulation and delivery via LNPs

The enhanced encapsulation and delivery observed at lower pH, as well as the need for an additional cationic lipid, initially indicated that electrostatic interactions played a role in LNP PepTAC encapsulation. However, the ^CR^PepTAC sequence carries a net charge of +2 at pH 7 and a theoretical isoelectric point of 9.8. At the formulation pH of 3, all acidic residues on the ^CR^PepTAC would be protonated, making it unlikely for encapsulation to solely rely on cation-anion interactions. Upon closer scrutiny of the sequence, we observed that the Ahx linker and E3 binding ligand display a strong hydrophobic sequence patch, making the ^CR^PepTAC sequence amphiphilic (Figure 2A). Based on this observation, we hypothesized that amphiphilicity plays a pivotal role in LNP ^CR^PepTAC encapsulation and delivery. To test this hypothesis, we prepared a fluorophore-labeled CREPT peptide-binding ligand (CL) without the linker and E3 domain and formulated it with the optimized LNP formulation. As depicted in Figure 4A, delivery was severely impaired with the CL alone. Encapsulation studies confirmed low encapsulation efficiency led to poor cellular uptake (Figure 4B). The data presented in Figures 4A and 4B strongly support the implication of peptide amphiphilicity as a crucial factor in PepTAC encapsulation and delivery. To showcase the generalizability of “amphiphilic-driven delivery”, we substituted the linker and E3 binding peptide with saturated alkyl chains of varying lengths (C6, C10, C12) in the ^CR^PepTAC. Delivery of these lipopeptide constructs in LNPs resulted in robust transfection, with longer lipid tails exhibiting improved transfection efficiency (Figure 4C, Supplementary Figure S17).

**Figure 4.**
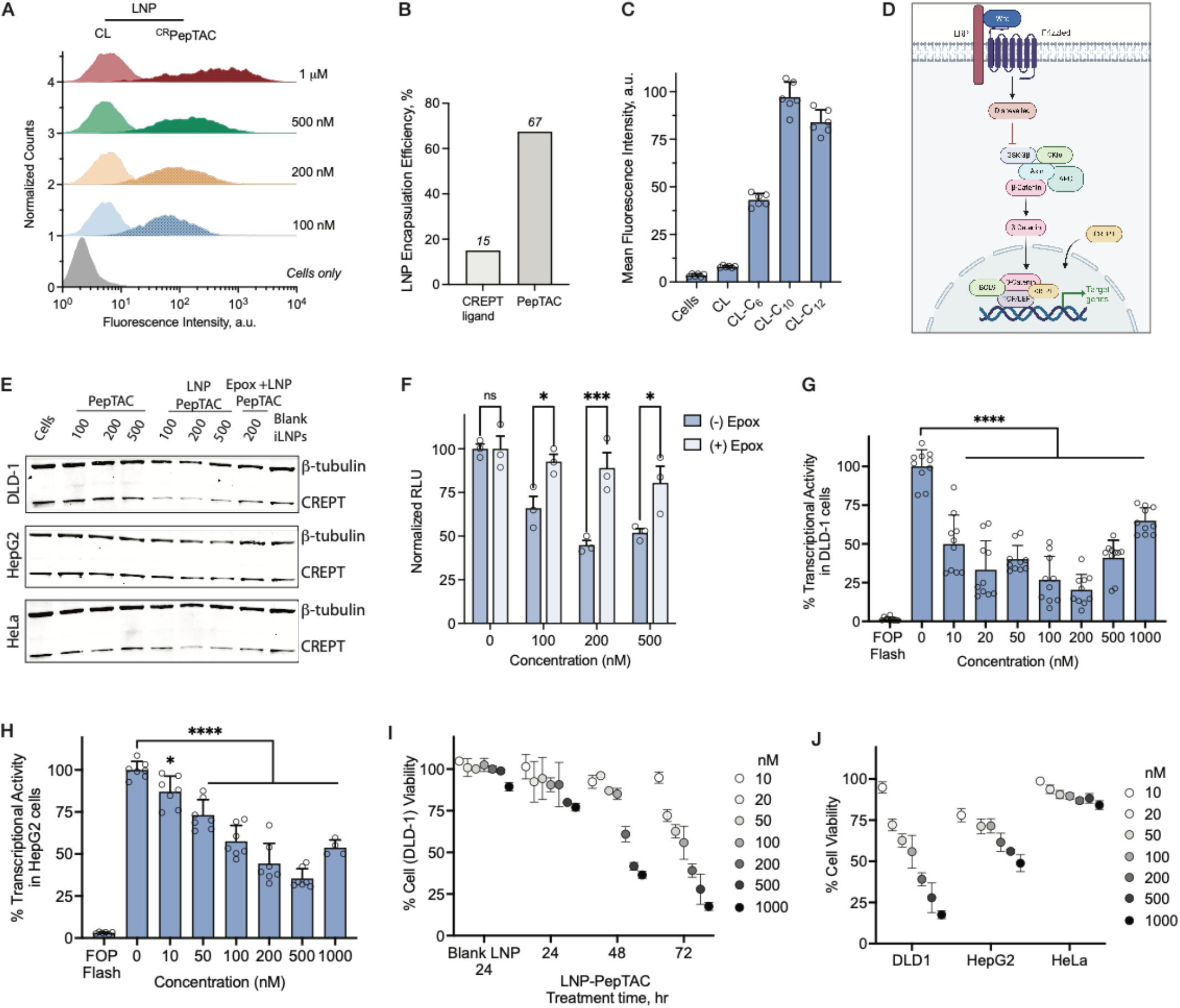
(A) HeLa cellular uptake of LNPs formulated with ^CR^PepTAC or peptide ligand alone at various concentrations. (B) Encapsulation efficiency of LNPs formulated with ^CR^PepTAC or peptide ligand alone. (C) HeLa cell uptake of CREPT ligand (CL) lipopeptides (500 nM) with different alkyl tail lengths. (D) Wnt signalling pathway and CREPT-initiated transcription. (E) Western blots of CREPT protein degradation in different cells with ^CR^PepTAC alone, formulated with iLNPs with and without the proteosome inhibitor, epoxymicin (Epox). Cells were treated for 24h and then harvested (F) Degradation of CREPT-Luc in HeLa cells treated with LNP-^CR^PepTAC for 24 h with and without epoxymicin (Epox). The data was normalized to Renilla expression in the same cells. Wnt active DLD-1 cells (G) and HepG2 cells (H) were transfected with Topflash and Fopflash luciferase followed by different concentrations of LNP-^CR^PepTAC. Luciferase reporter activity was assayed 24 hours after ^CR^PepTAC treatment. (I) Effect of LNP-^CR^PepTAC treatments at different concentrations on Wnt-active DLD-1 cell viability at 24, 48 and 72h. Cell viability was assessed via an MTS assay. (J) Effect of LNP-^CR^PepTAC treatments at different concentrations on Wnt-active DLD-1 and HepG2, and non-Wnt active HeLa cell viability after 72h. Cell viability was assessed via an MTS assay. Data are displayed as mean ± SD by one-way ANOVA. * p < 0.05, ** p < 0.01, ***p < 0.001, ****p < 0.0001.

### ^CR^PepTACs delivered by LNPs enable CREPT degradation

After establishing that LNPs enhance the cellular uptake of PepTACs, we next focused our investigations on assessing the protein degradation activity of the delivered ^CR^PepTAC. CREPT is a highly tumorigenic protein overexpressed in Wnt-activated malignant cells and tissues. It functions as a crucial regulator of genes such as *CCND1* that enhance cell proliferation and promote tumorigenesis (Figure 4D).^41,42^ Considering CREPT’s role in transcriptional activation, intracellular degradation of CREPT (i.e. RPRD1B) would disrupt the Wnt signaling pathway, directly impacting cell proliferation. To assess the functional delivery and efficacy of PepTAC-LNP formulations, we assayed the endogenous expression of CREPT in cells following ^CR^PepTAC-LNP treatment. Our results demonstrated robust degradation of endogenous CREPT protein across various clinically relevant cell lines (Figure 4E). Treatment with epoxomicin, a proteasome inhibitor, prevented protein degradation underscoring the role of the UPS in ^CR^PepTAC-mediated protein degradation. Furthermore, we quantitatively evaluated CREPT protein degradation using a firefly luciferase-fused CREPT (Luc-CREPT) expressing plasmid. Cells were pre-transfected with Luc-CREPT and a Renilla luciferase as an internal control, then treated with varying amounts of ^CR^PepTAC-LNP formulations. We observed dose-dependent degradation of Luc-CREPT protein (Figure 4F). Again, degradation was inhibited in the presence of the epoxomicin. Degradation of Luc-CREPT slightly decreased at high concentrations (>200 nM) likely due to the “hook” effect, a phenomenon observed in many PROTACs. This occurs when PepTACs (including PROTACs) saturate binding to their protein target and E3 ligase, leading to the formation of binary complexes instead of the ternary complex required for ubiquitination and degradation.^45-47^ We observed 50% degradation (DC^50^) of CREPT at concentrations between 100-200 nM, representing one of the lowest reported DC^50^ among all PepTACs to the best of our knowledge. Finally, we assayed for CREPT degradation with and without DOTAP and observed diminished degradation when DOTAP was absent (Supplementary Figure S18).

We next carried out a TopFlash assay using two constitutive Wnt-activated cell lines, DLD-1 and HepG2, to assess whether ^CR^PepTAC-LNP-mediated CREPT degradation could suppress Wnt signaling and transcriptional activity. The TopFlash construct is a T cell factor (TCF)/lymphoid enhancer-binding factor (LEF)-Firefly luciferase reporter vector that is activated in Wnt-active cells.^48,49^ The firefly luciferase gene in this reporter is controlled by the TCF/LEF responsive element, which is activated in the presence of CREPT, β-catenin and other transcriptional proteins. Therefore, degradation of CREPT should lead to a decrease in transcriptional activity and luciferase expression. As a control, we used a non-inducible luciferase vector called FopFlash, which is under the control of a minimal promoter without any response elements, thus providing a measure of background luciferase activity. In both Wnt-activated DLD-1 and HepG2 cells, we observed a significant concentration-dependent reduction in transcriptional activity following ^CR^PepTAC-LNP treatment (Figure 4G and 4H). In DLD-1 cells, a 50% suppression of transcriptional activity was observed at concentrations as low as 10 nM (Figure 4G). Consistent with previous observations, we observed the intrinsic “hook” effect at high concentrations, occurring beyond 200 nM in DLD-1 cells and beyond 500 nM in HepG2 cells (Figure 4G and 4H).

A decrease in the transcription of proliferative genes in Wnt-active cell lines should result in the suppression of cell growth. To assess this downstream effect, we conducted a time-dependent MTS assay to examine the impact of the ^CR^PepTAC-LNP formulation on cell proliferation and viability in Wnt-active DLD-1 and HepG2 cells, as well as non-Wnt-active HeLa cells. We observed a significant reduction in cell proliferation over time in Wnt-active DLD-1 (Figure 4I) and HepG2 cells (Figure S19). This decrease in cell viability was specific to Wnt-active DLD-1 and HepG2 cells, as we observed minimal changes in the viability of non-Wnt-active HeLa cells 72 hours after treatment (Figure 4J). These collective results demonstrate the ability of ^CR^PepTAC-LNP to deliver PepTACs into the cytosol to degrade the target protein, CREPT. Protein degradation leads to a loss of function, as evidenced by the concurrent decrease in transcriptional activity and cell proliferation.

### ^βCat^PepTACs delivered by LNPs enable robust β-catenin degradation

To demonstrate the versatility of the LNP-mediated PepTAC delivery strategy, we designed a novel PepTAC targeting a well-studied oncoprotein, β-catenin. The design was based on a peptide derived from the BCL9 protein that binds to β-catenin as seen in the co-crystal structure^50-52^ and reproduced by AlphaFold (Figure 5A). β-catenin was selected given its crucial role in the Wnt-signaling pathway of tissue homeostasis and embryonic development, as well as in several types of human cancer, such as colorectal, breast, melanoma, and prostate, among several others.^53-56^ Like CREPT, β-catenin controls the expression of several key genes that regulate cell cycle, proliferation and tumorigenesis. Transcriptional activation of the Wnt/β-catenin pathway is dependent on formation of the β-catenin super complex involving BCL9 and TCF/LEF family of transcriptional factors. As such, molecules that degrade β-catenin can inhibit Wnt/β-catenin signal transduction and supress cell proliferation.^50,51,53,55,57,58^ To construct a PepTAC against β-catenin (^βCat^PepTAC), we introduced the Ahx linker and the pentapeptide VHL-binder at the N-terminal of the β-catenin peptide binding sequence (SQEQLEHRERSLQTLRDIQRMLF). This N-terminal region is solvent exposed (Figure 5A) and is expected to not interfere with ligand binding to its target.

**Figure 5.**
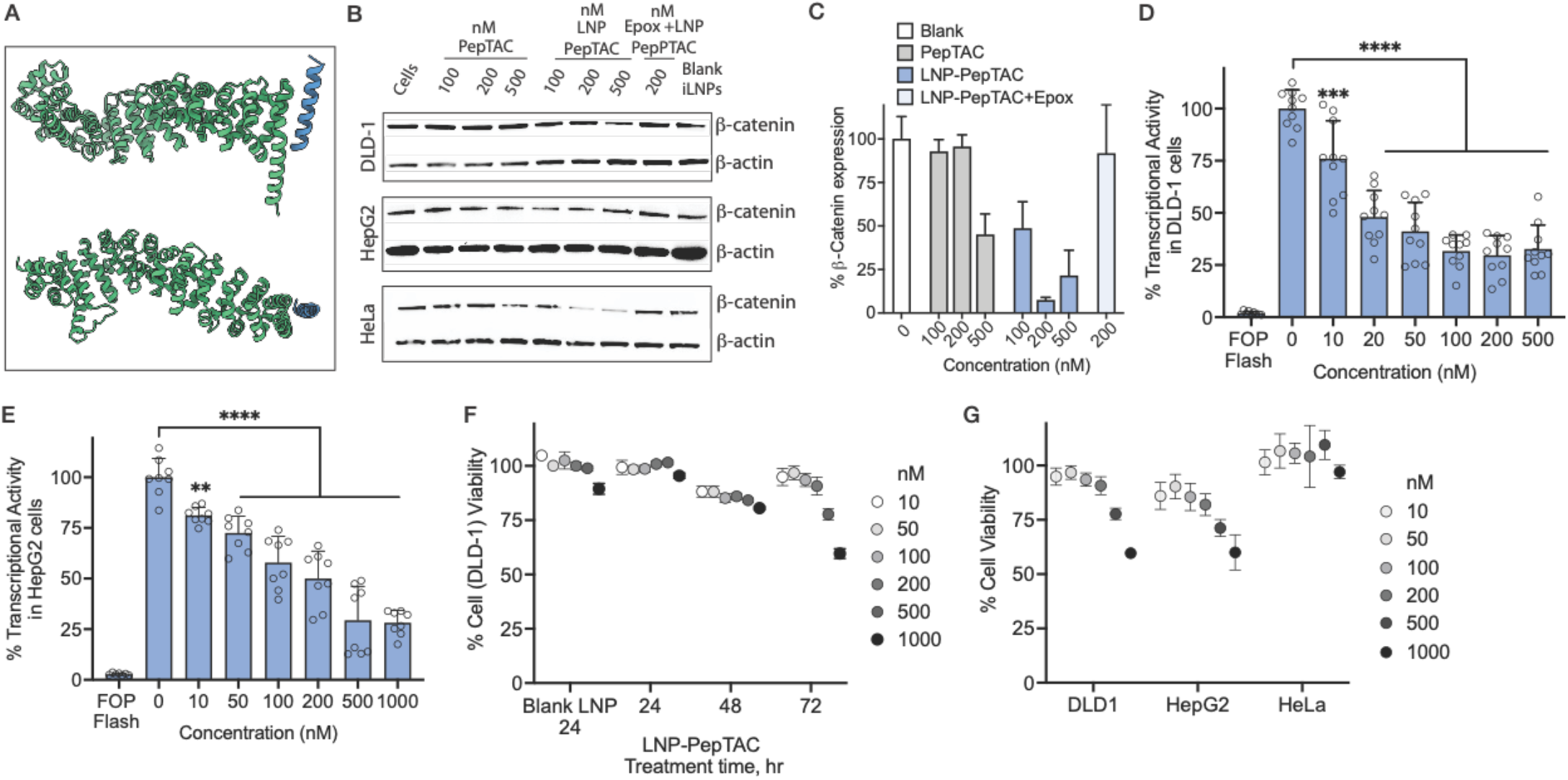
(A) AlphaFold-Multimer prediction of β-catenin (armadillo repeat region) with the BCL9 derived peptide ligand. (B) Western blots of β-catenin protein degradation in different cells with ^βCat^PepTAC alone, formulated with LNPs with and without the proteosome inhibitor, epoxymicin. Cells were treated for 24h and then harvested. (C) Quantification of HeLa cell western blot (duplicates). Wnt active DLD-1 cells (D) and HepG2 cells (E) were transfected with Topflash and Fopflash luciferase followed by different concentrations of LNP-^βCat^PepTAC. Luciferase reporter activity was assayed 24 hours after ^βCat^PepTAC treatment. (F) Effect of LNP-^βCat^PepTAC treatments at different concentrations on Wnt-active DLD-1 cell viability at 24, 48 and 72h. Cell viability was assessed via an MTS assay. (G) Effect of LNP-^βCat^PepTAC treatments at different concentrations on Wnt-active DLD-1 and HepG2, and non-Wnt active HeLa cell viability after 72h. Cell viability was assessed via an MTS assay. Data are displayed as mean ± SD by one-way ANOVA. * * p < 0.05, ** p < 0.01, ***p < 0.001, ****p < 0.0001.

Functional LNP-mediated delivery of ^βCat^PepTAC was assessed by measuring the endogenous cellular expression levels of β-catenin. Our results demonstrated robust degradation of β-catenin in clinically relevant cell lines (Figure 5B). β-catenin degradation was inhibited by treatment with the proteasome inhibitor epoxomicin, indicating degradation through the UPS (Figure 5B and 5C). We next examined downstream effects of β-catenin degradation by conducting a TopFlash assay using two constitutive Wnt-activated cell lines, DLD-1 and HepG2. Degradation of β-catenin led to a significant concentration-dependent reduction in transcriptional activity following ^βCat^PepTAC-LNP treatment (Figure 5D and 5E). A 50% suppression of transcriptional activity was observed at concentrations as low as 20 nM in DLD-1 cells (Figure 5D) and at 100 nM in HepG2 cells (Figure 5E). The intrinsic “hook” effect isn’t observed in either of the Wnt-active cells and likely exists at concentrations higher than those tested.

Finally, the downstream effect of transcriptional suppression was assessed by performing time-dependent cell viability studies on Wnt-active DLD-1 and HepG2 cells, as well as non-Wnt-active HeLa cells. Similar to the studies with CREPT, we observed a concentration-dependent reduction in cell viability over time in Wnt-active DLD-1 (Figure 5F) and HepG2 cells (Figure S20). The effect on cell viability was specific to Wnt-active cells, as we observed minimal changes in the viability of non-Wnt-active HeLa cells 72 hours after treatment (Figure 5G) despite evidence of β-catenin degradation in these cells (Figure 5B and 5C). These collective results demonstrate the delivery of ^βCat^PepTACs by LNPs to the cytosol, enabling the targeted degradation of β-catenin. β-catenin degradation, like CREPT, led to a decrease in transcriptional activity and cell proliferation.

### LNPs extend PepTAC half-life in vivo, enable biodistribution to the liver and spleen, and facilitate β-catenin degradation in the liver

The current results demonstrate that LNPs facilitate PepTAC delivery into cells and that the delivered PepTACs retain their protein-degrading capabilities. To further explore their suitability for potential in vivo applications, we investigated the distribution of PepTAC-LNPs upon systemic administration. Recent studies highlight the significance of the apparent surface pKa of LNPs as a critical factor influencing the protein corona composition and LNP biodistribution upon systemic administration.^59,60^ Notably, LNPs with apparent pKa values ranging from 6 to 7 have been observed to accumulate primarily in the liver.^59^ Considering this link between pKa and biodistribution, we determined the apparent pKa of PepTAC-LNPs to be 6.5 using the 6-(p-toluidino)-2-naphthalenesulfonic acid (TNS) assay indicating preferential accumulation in the liver (Supplementary Figure S21).^67^ Indeed, biodistribution experiments in hairless SKH-1 mice using Cy5.5-labeled PepTAC-LNPs demonstrated a prominent accumulation of PepTAC-LNPs in the region of the liver (Figure 6A and 6B). Peak accumulation was observed at 1 hour post-injection (Figure 6A and 6B). Quantitative analyses demonstrated that LNPs significantly increased PepTAC accumulation in various organs, with a notable enhancement in the liver and spleen, compared to PepTAC alone (Figure 6B and 6C). Given their preferential distribution to the liver, we assessed cellular penetration of PepTAC-LNPs into liver cells. Confocal imaging of liver sections revealed significant intracellular accumulation of PepTAC-LNPs into liver cells at the 1 hour time point relative to PepTAC alone. After 24 hours, PepTAC signal from PepTAC-LNPs are still visible in the liver sections where no signal is observed with PepTACs alone. This data demonstrates that LNPs extend PepTAC accumulation in vivo relative to PepTAC alone.

**Figure 6.**
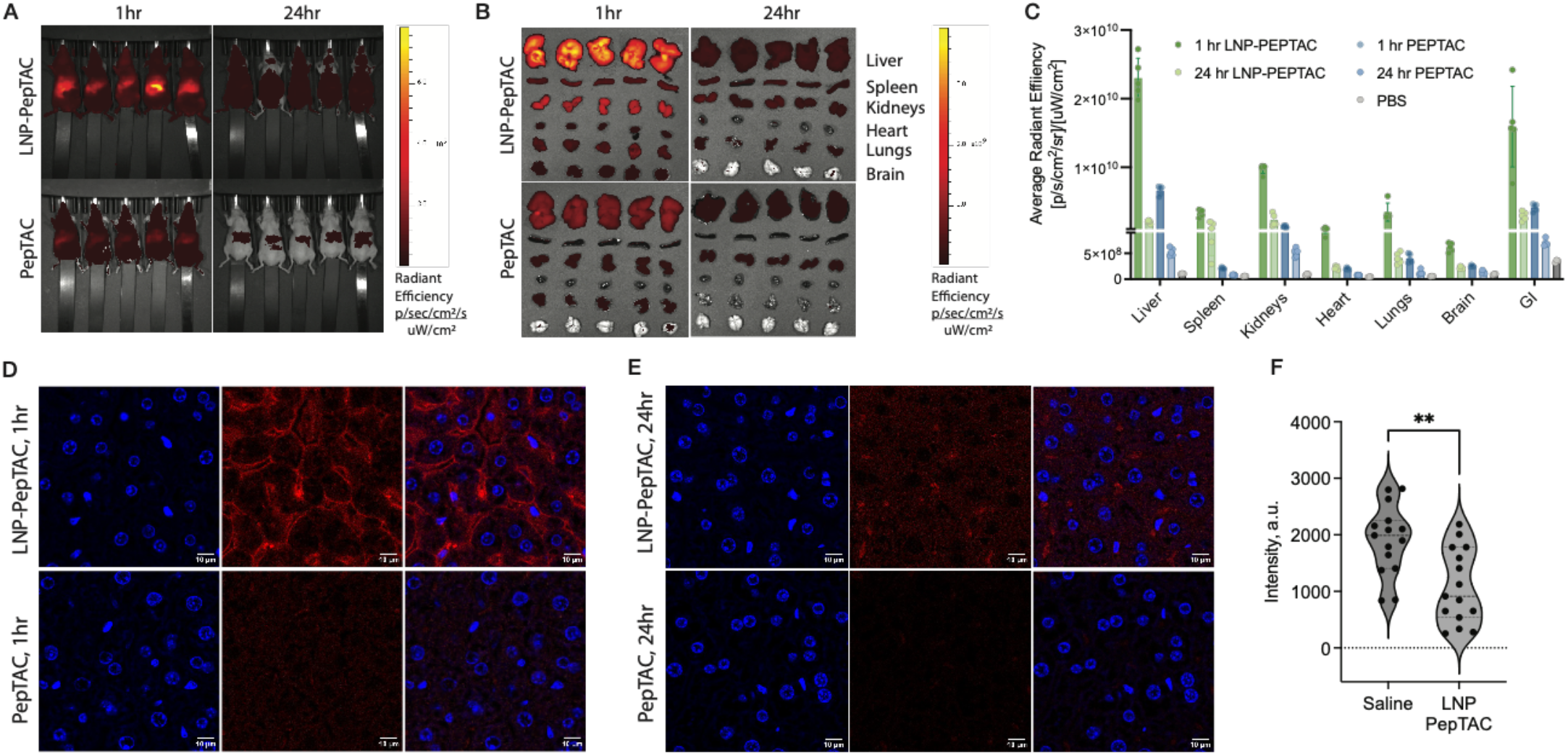
(A) Whole animal fluorescence imaging of SKH-1 hairless mice after 1 hour and 24 hours. Top - LNP-^CR^PepTAC, bottom ^CR^PepTAC only (5 mg/kg, 50% labeled cargo). Ex vivo imaging (B) and quantified fluorescence signal (C) after 1 hour and 24 hours of Cy5.5 labeled PepTACs in major organs extracted from SKH-1 hairless mice IV injected with LNP-PepTAC, PepTAC alone and PBS. Confocal images of sectioned liver tissue that received Cy5.5 labeled LNP-PepTAC or PepTAC alone after 1 hour (D) and 24 hours (E) stained with nuclei stain (Hoechst, left panel). Cy5.5 fluorescence represented as a false red color (middle panel) and the overlay is shown on the right panel. (F) Liver β-catenin levels assessed via western blot from normal healthy Balb/c mice treated every 3 days for 10 days with saline or 5 mg/kg ^βCat^PepTAC-LNPs. Data are displayed as a violin plot and analyzed using an unpaired t-test with Welch’s correction. * p < 0.05, ** p < 0.01, ***p < 0.001, ****p < 0.0001.

Given their preferential distribution to the liver, we assessed the ability of the ^βCat^PepTACs encapsulated in LNPs to downregulate liver β-catenin in wild type mice. Mice were intravenously injected every 72 hours for ten days (four injections) with ^βCat^PepTAC-LNPs. Following this treatment, liver tissue samples were harvested, digested, and analyzed for presence of β-catenin via Western Blot.

β-catenin levels were significantly reduced in mice treated with ^βCat^PepTAC-LNPs relative to saline control (Figure 6F, Supplementary Figure S22). In healthy mice, the expression of β-catenin in liver hepatocytes is relatively low, primarily due to the absence of Wnt signalling, which leads to its rapid endogenous degradation.^61^ Considering its natural low expression, the detection of ∼50% β-catenin degradation following ^βCat^PepTAC-LNP treatment is noteworthy and holds promise for future applications, especially in disease states characterized by the overexpression of β-catenin due to Wnt pathway activation.

## DISCUSSION

Poor cellular permeability and inherent instability in biological fluids are major obstacles that have impeded peptide-based drug development. To overcome these barriers, we repurposed the LNP formulation to enable efficient intracellular PepTAC delivery at nanomolar concentrations. LNPs can shield the PepTAC cargo from extracellular proteases, promote cellular uptake, facilitate PepTAC escape from endosomes, and enable PepTAC-mediated ubiquitination of target proteins followed by proteosome degradation. Unlike nucleic acids which load via electrostatic-driven assembly into LNPs, we show PepTAC amphiphilicity as the key feature that enables encapsulation and delivery by LNPs (Figure 4).

We formulated PepTACs against two transcription factors involved in the Wnt signaling pathway: CREPT and β-catenin, which is notably difficult to drug. Our results showed significant degradation of CREPT and β-catenin across various clinically relevant cell lines at concentrations as low as 100 nM, which is unprecedented for PepTACs. Critically, neither ^CR^PepTAC alone nor ^CR^PepTAC modified with a cell-penetrating peptide sequence induced degradation at nanomolar concentrations, (Figure 4E) which is consistent with multiple reports that show the onset of PepTAC-mediated degradation in micromolar range.^24,26,27,30-33,62,63^ Similar results were achieved with ^βCat^PepTAC-LNP formulations (Figure 5) with robust β-catenin degradation at nanomolar ^βCat^PepTAC concentrations. Considering their role in transcriptional activation, intracellular degradation of CREPT and β-catenin disrupted the Wnt-dependent transcriptional activity and directly impacted cell proliferation in Wnt-dependent cancer cells (DLD-1 and HepG2), with no discernible effect on Wnt-independent cells (HeLa).

Finally, we demonstrate the in vivo potential of these formulations by illustrating enhanced accumulation of LNP-delivered PepTACs in the liver and spleen and significant β-catenin degradation in the liver. Our current demonstration was limited to wild-type mice, as we aimed to de-risk this novel approach and establish the feasibility of PepTAC delivery in vivo. One potential application of this delivery system is in the development of therapeutics for hepatocellular carcinoma (HCC), given that hMet and β-catenin mutations account for approximately 10% of all HCC cases.^64-67^ Future studies will involve employing an HCC model with *CTNNB1* mutations to assess the efficacy of the ^CR^PepTAC- and ^βCat^PepTAC-LNP formulations as therapeutics. As LNP technology evolves for extrahepatic targeting, we anticipate that, akin to siRNA and other nucleic acid technologies, PepTACs will emerge as a modality for manipulating protein levels in various organs. Unlike siRNAs, PepTACs can induce rapid direct protein degradation, irrespective of protein half-life, and can recognize different protein states, such as mutant vs. wild type or active vs. inactive conformation, allowing for the depletion of specific protein subpopulations.^53,68,69^

In cases where life-saving medicines are urgently needed (e.g., to contain an outbreak), drug discovery strategies that can be rapidly developed from existing protein structural data and widely deployed are of paramount importance. By overcoming the intracellular transport limitations of PepTACs, we have expanded their use broadly as novel chemical biology tools for dissecting protein networks and as potential therapeutic modalities for inhibiting new drug targets that have so far evaded pharmacological intervention. Our vision is to usher in a new era in which PepTACs can be readily procured, much like the manner in which siRNAs are currently procured against specific mRNA targets. The continuous advancement of computational techniques, exemplified by AlphaFold and Rosetta Peptiderive,^23^ is rapidly propelling us toward the ability to computationally predict peptide binding sequences for any known protein target with established protein-protein interactions. As this capability becomes a reality, this study will serve as a guide for optimizing intracellular delivery of PepTACs. This optimization will, in turn, facilitate widespread adoption of PepTACs for both basic research and therapeutic applications.

### Online content

Any methods, additional references, source data, extended data, supplementary information, acknowledgements, peer review information; details of author contributions and competing interests; and statements of data and code availability are available at: xxx.

## METHODS

### Peptide synthesis

All peptide sequences were synthesized using Rink Amide MBHA resin (100-200 mesh) with an average loading efficiency of 0.93 mmol/g, FMOC-protected amino acids, DMF as the solvent, Diisopropylcarbodiimide (DIC) as the activator and oxyma as the activator base. Solid-phase peptide synthesis was carried out in a microwave-assisted automated solid-phase peptide synthesizer at a 0.1 mmol synthesis scale. The following peptide sequences were synthesized:

CREPT ligand (CL): KDVLSEKEKKLEEYKQKLARV

^CR^PepTAC: IYP^OH^AL-Ahx-KDVLSEKEKKLEEYKQKLARV

^βCat^PepTAC: IYP^OH^AL-Ahx-SQEQLEHRERSLQTLRDIQRMLF

P^OH^ stands for hydroxylated proline and Ahx is 6-aminohexanoic acid. All sequences are written from N-terminal to C-terminal. Following peptide synthesis, the resins were cleaved using a cleavage cocktail consisting of 92.5% trifluoroacetic acid (TFA), 2.5% water, 2.5% triisopropylsilane (TIPS) and 2.5% phenol. Cold diethyl ether was added to the cleaved crude mixture which led to peptide precipitation as a white powder. Crude peptide was purified via reverse-phase HPLC (using water/acetonitrile with 0.1% trifluoroacetic acid as the solvent system and a 5-100% acetonitrile gradient in 20 minutes). The purified peptide fraction was characterized using MALDI-TOF mass spectroscopy and the purity was confirmed using an Agilent 1100 analytical HPLC.

### Non-specific peptide labeling with Fluorescein-NHS

Purified peptides were dissolved in DMSO to a concentration of 25 mg/mL. Fluorescein-NHS ester (5/6-carboxyfluorescein succinimidyl ester, 1.1 molar equivalents) and diisopropylethyl amine (DIPEA) (2.5 molar equivalents) were then combined and subsequently added to the peptide solution. The reaction mixture was stirred at room temperature for 2 hours and purified by RP-HPLC (using water/acetonitrile with 0.1% trifluoroacetic acid as the solvent system and a 5-100% acetonitrile gradient in 20 minutes). The purified labeled peptides were lyophilized and characterized by MALDI-TOF MS.

### N-terminal labeling of peptides with fluorophore-NHS esters

The immobilized ^CR^PepTAC peptide on a rink-amide resin (20mg) with a free N-terminal, all other amino acid side-chains protected, was swollen in 0.5 mL of DMF for 10 minutes. 0.4 mL of DMF was aspirated and 0.1 mL DMF with either fluorescein-NHS ester or Cy 5.5-NHS ester (5 molar equivalents) and DIPEA (10 molar equivalents) were added. The reaction mixture was stirred at room temperature for 12 hours. Subsequently, the resin was washed overnight with DMF and finally cleaved with 92.5% TFA, 2.5% water, 2.5% TIPS, and 2.5% phenol. The cleaved crude labeled peptide was dried with a stream of nitrogen, precipitated with the addition of pre-cooled ether, re-dissolved in water/acetonitrile (1:1 volume ratio), purified using RP-HPLC and characterized via MALDI-TOF MS.

### N-terminal lipidation and fluorescein modification of peptides

The immobilized CL peptide on a rink-amide resin (20mg) with a free N-terminal, all other amino acid side-chains protected, was swollen in 0.5 mL of DMF for 10 minutes. 0.4 mL of DMF was aspirated. Solutions of hexanoic, decanoic and dodecanoic acid (2.5 molar equivalents) were pre-incubated with HATU (2.5 molar equivalent) and DMAP (3 molar equivalents) dissolved in 0.1 mL DMF. The solutions were added to the swelled resin and stirred at room temperature for 12 hours. Subsequently, the resin was washed 3-5 times with both DMF and diethyl, and dried overnight. Cleavage from the resin was done with 92.5% TFA, 2.5% water, 2.5% TIPS, and 2.5% phenol. The cleaved, crude, 8lipid-modified CL-peptide was dried with a stream of nitrogen and precipitated via cold-ether precipitation at -80°C. The 8lipid-modified peptides were re-dissolved in water/acetonitrile (1:1 volume ratio), and purified using RP-HPLC followed by MALDI-TOF for characterization. Purified peptides were non-specifically labelled with fluorescein dye as previously described.

### LNP Formulation for in vitro experiments

To make 100 µL of LNP formulation with 25 µg of peptides as the payload, D-Lin-MC3 (45 mol%, 5 µL of 10 mg/mL ethanol stock), DSPC (8.9 mol%, 6.15 µL of 2 mg/mL ethanol stock), cholesterol (34.7 mol%, 4.64 µL of 5 mg/mL ethanol stock), DMG-PEG-2000 (1.4 mol%, 2.93 µL of 2 mg/mL ethanol stock) and DOTAP (10 mol%, 2.42 µL of 5 mg/mL ethanol stock) were mixed with 3.9 µL of ethanol to make a 25 µL lipid-cocktail. Subsequently, 5 µL of 5 mg/mL (25 µg) PepTAC was diluted with 20 µL of 10 mM sodium citrate buffer solution (pH 3) to obtain a total of 25 µL aqueous PepTAC-solution. A micropipette was used to rapidly mix the lipid-cocktail with the aqueous PepTAC-solution. This mixture was then diluted with an equal volume of 1xPBS buffer (pH 7.4, 50 µL) to make a formulation with a pre-dialysis volume 100 µL. This formulation was then dialyzed in a dialysis chamber (Slide-A-Lyzer™ MINI Dialysis Devices, 3.5K MWCO, Thermo Fisher) against sterile 1x PBS, pH 7.4 for 2 hours. The formulation was stored at 4 ^0^C.

To create LNP formulations at different pH values, a sodium citrate buffer of different pH values was used to prepare the 25 µL PepTAC-solution. Furthermore, 10 mol% of DDAB, DODAP, or EPC lipids were used in place of DOTAP for making formulations with varying compositions of the 5^th^ lipid. Similarly, varying molar percentages of DOTAP supplements (5-30%) were used to create different LNP formulations.

### LNP formulation for in vivo experiments

To make 100 µL of LNP formulation with 100 µg of peptides as the payload, D-Lin-MC3 (45 mol%, 8 µL of 25 mg/mL ethanol stock), DSPC (8.9 mol%, 6.56 µL of 7.5 mg/mL ethanol stock), cholesterol (34.7 mol%, 4.64 µL of 20 mg/mL ethanol stock), DMG-PEG-2000 (1.4 mol%, 2.34 µL of 10 mg/mL ethanol stock) and DOTAP (10 mol%, 1.93 µL of 25 mg/mL ethanol stock) were mixed with 3.9 µL of ethanol to make a 25 µL lipid-cocktail. Next, 4 µL of 25 mg/mL (100 µg) of PepTAC was added to 21 µL of 10 mM sodium citrate buffer solution (pH 3) to make a total of 25 µL of aqueous PepTAC-solution. For biodistribution experiments, 25 µL of aqueous PepTAC-solution was made by mixing 19.4 µL of sodium citrate buffer, 2 µL of 5 mg/mL Cy 5.5-labeled ^CR^PepTAC (to acheive 10% labeling) and 3.6 µL of 25 mg/mL unlabeled ^CR^PepTAC peptide. A micropipette was used to rapidly mix the lipid-cocktail with the aqueous PepTAC-solution. This mixture was then rapidly diluted with an equal volume of 1X PBS buffer (pH 7.4, 50 µL) to make a formulation with a pre-dialysis volume 100 µL. The formulation was dialyzed in a dialysis chamber (Slide-A-Lyzer™ MINI Dialysis Devices, 3.5K MWCO, Thermo Fisher) against sterile 1X PBS, pH 7.4 for 4 hours. The formulation was stored at 4^0^C.

### Size and zeta potential measurements

PepTAC-LNP formulations were diluted 50-fold in 1X PBS buffer. A total of 500 µL of the diluted formulations were placed in a polystyrene cuvette with a 10 mm path length and particle size measurements were made with a Malvern Zetasizer. The same formulation was placed in a Malvern capillary sample cell for zeta potential measurements.

### Apparent pKa measurement via a TNS assay

A series of buffers with pH values between 2.5 and 8.5 were prepared by adjusting the pH of a solution containing 10 mM citrate, 10 mM phosphate, 10 mM borate, and 150 mM NaCl with 1 N HCl and 5 N NaOH. 80 µL/well of each buffer solution was added to a black-wall 96-well plate. A 300 µM stock solution of TNS was prepared in DMSO and 2 µL of this solution was added to each of the buffer solutions in the 96-well plate. 20 µL of a particular LNP formulation was added to each well (with replicates). The fluorescence of the resulting solution was measured using a microplate reader (TECAN Infinity M1000 Pro) using an excitation wavelength of 325 nm and an emission wavelength of 435 nm with 10 nm bandwidth. The emission intensity was plotted against pH and fit using a four-parameter sigmoidal logistic equation (GraphPad Prism) to obtain the apparent LNP surface pKa.

### Encapsulation efficiency of the LNPs

15 µL of each LNP formulation (0.25 mg/ml with respect to labelled peptide) was placed in an Eppendorf tube and mixed with 5 µL of 5X Triton-X solution prepared with in PBS buffer followed by sonication for 10 minutes to lyse the particles. In a separate Eppendorf, another 15 µL of the same formulation was diluted with an equivalent 5 µL of 1X PBS buffer. Both lysed and intact LNP solutions were mixed with 20 µL of the native gel loading buffer and loaded in a 16% Tricine gel for a native gel electrophoresis. In-gel fluorescence of the non-encapsulated labeled payload from the lysed and intact LNP solutions was measured and quantified via ImageJ. Quantification using the relationship; [(*l*_*lys*e*d* ™ *l*_i*ntact*)/*I*_*lys*e*d*] × 100, gives a measurement of the % encapsulation efficiency.

### Cloning for Firefly Luciferase-fused CREPT/RPRD1B expressing plasmid

Construction of pCMV-Luc-CREPT was performed using restriction cloning and propagated in Escherichia coli strain DH5*a*. Firefly luciferase was PCR amplified from pCDNA-Luciferase (Addgene, cat # 18964) using primers introducing a 5’ KpnI site and a 3’ NheI site, and CREPT was PRC amplified from pCMV-RPRD1 (Sino Biological, cat # HG14027-NH) using primers introducing a 5’ NheI site and a 3’ XbaI site. The two gene fragments were ligated using NheI and then ligated into pCMV (Sino Biological) using KpnI and NheI. The primer sequences are given below:

Luc forward primer: 5’-GGTACCATGGAAGACGCCAAAAACATAAAGAAAGG-3’ Luc reverse primer: 5’-GCTAGCGCCTCCACCCACGGCGATCTTTCCGCC-3’ CREPT forward primer: 5’-GCTAGCTCCTCCTTCTCTGAGTCGGC-3’ CREPT reverse primer: 5’-TCTAGATTAATGATGGTGGTGATGGTGGTCAGTTGAAAACAGGTCCCCAG-3’

### Cell culture, imaging and cellular assays

HeLa and HepG2 cells were grown in DMEM media supplemented with 10% FBS (fetal bovine serum) and 1% penicillin/streptomycin, while DLD1-cells were grown in RPMI 1600 media with the same supplement. SK-OV-3 and SK-BR-3 cells were grown in McCoy 5A media with the same supplement. During cell-passaging, trypsin-EDTA (0.25%) with phenol red was used for the detachment of cells from the tissue-culture flasks. All cells were maintained at 37 ^0^C, 5% CO_2_ and 90% relative humidity.

### Cell-uptake assay via flow cytometry

75,000 cells were seeded per well in 500 µL of DMEM media supplemented with 10% FBS and 1% penicillin/streptomycin in a 24 well-plate. The plate was incubated for 18-20 hours at 37 ^0^C with 5% CO_2_. Free fluorescein-labeled peptides and LNP-encapsulated fluorescein-labeled peptides in PBS buffer were added at various concentrations to the wells. In all LNP formulations, the PepTAC dose added to cells was based on the total amount of PepTAC added to the formulation. Thus, a PepTAC concentration of 0.25 mg/mL was used for the 100 uL formulations. After treatment, the plate was incubated for another 20-24 hours (kinetic studies were incubated for different time points). The media was aspirated, and cells were washed three times with 1x PBS. Cells were then detached by pipetting 20-25 times with PBS. Detached cells were centrifuged (1200 x g for 6 mins), the supernatant aspirated, and pelleted cells were resuspended in fresh 1X PBS for flow cytometry studies. Readings were taken on a FACSCalibur™ analyzer (Becton Dickinson) with fixed power and gain parameters. Flow cytometry data were analyzed using the FlowJo software. The fluorescence histogram and mean fluorescence intensity (MFI) data were obtained from 10,000-gated cells.

### Confocal imaging

70,000 HeLa cells/chamber were plated in a 4-chamber 35-mm glass-bottom microwell dish (MatTeK) and cultured at 37^0^C for 20-24 hrs with 500 µL of DMEM media supplemented with 10% FBS and 1% penicillin/streptomycin. Each chamber was treated with different LNP formulations containing fluorescein-labeled peptides or free labeled peptides as controls. The chambers were incubated for 8 hours. Thereafter, media was aspirated, and each chamber was washed 2-3 times with 1X sterile PBS. The chamber was then filled with 500 µL of dye-free Fluorbrite DMEM media pre-mixed with 1x Hoechst (1:12000 dilution of Thermo Hoechst 33342), 1X endosome-tracker 594 (1:1000 dilution of Biotium LysoView™ 594) and 1X microtubule-tracker 647 (1:1000 dilution of Biotium ViaFluor^R^ Live cell microtubule stain 647). After incubation for 10-15 minutes, the media was aspirated, and each chamber was washed 3-4 times with 1X PBS and re-filled with dye-free Fluorbrite DMEM media. The chamber was then subjected to live cell imaging on a Zeiss i880 inverted confocal microscope consisting of four LASER channels; the blue channel (405 nm excitation) to probe the Hoechst nuclei stain, the green channel (488 nm excitation) to probe the labeled PepTAC, the red channel (594 nm excitation) to probe endosomes and the far-red channel (647 nm, represented as grayscale) to probe cytosolic microtubules. The comparative image at different channels and the merged image qualitatively assesses the sub-cellular localization and distribution of the delivered PepTAC cargo.

### Assessing intracellular target protein degradation via Western Blot

150,000-200,000 cells were plated per well of a 12-well plate with 1 mL of DMEM media. The plate was then incubated for 18-20 hours. Free peptides or LNP-encapsulated peptides were subsequently added to individual wells in 1X PBS buffer at various doses. In all LNP formulations, the PepTAC dose added to cells was based on the total amount of PepTAC added to the formulation. Thus, a PepTAC concentration of 0.25 mg/mL was used for the 100 µL formulations. After treatment, the plate was incubated for an additional 24 hours. Next, media was aspirated, and 75 µL of Pierce RIPA (Radio Immuno Precipitation Assay) lysis buffer, pre-mixed with 1X Halt protease inhibitor cocktail and 1X EDTA solution (5 mM final concentration), was added to each well at 0°C (on an ice bath) and incubated for 10-15 minutes with occasional stirring. Cell lysates were collected in pre-cooled sterile Eppendorf tubes, and the water-insoluble/membrane-soluble fractions were separated by centrifuging the samples at 20,000 x g for 15 minutes. The supernatants were collected, and 15 µL of each lysate sample was used for total protein quantification using a standard BCA assay (5 µL per measurement, with three replicates).

In the CREPT and β-catenin knockdown experiment, 20 µg and 15 µg of cell lysates respectively were loaded into each well of a 4-12% bis-tris Nu-PAGE gel and 1X MOPS running buffer was used to run the gels. Upon completion, the gel was transferred to a nitrocellulose membrane with a porosity of 0.45 µm using a wet-transfer apparatus and transfer buffer containing 1X Tris-glycine with 20% methanol. The membrane was then treated with a blocking buffer consisting of 1X TBST buffer with 5% milk powder for 2-3 hours. Following that treatment, the membrane was incubated overnight at 4°C with the primary antibody solution, which was prepared by diluting the rabbit anti-CREPT antibody (1:2500 dilution), mouse anti-β-Catenin antibody (1:1000 dilution), rabbit anti-β-tubulin antibody (1:2500 dilution), and mouse anti-β-actin antibody (1:2500 dilution) in the blocking buffer. The membrane was then washed 5-6 times for 5 minutes with 1X TBST buffer. Next, the membrane was treated with the secondary antibody solution, prepared by diluting the Starbright 700 Goat anti-rabbit IgG antibody (1:5000 dilution) and Starbright 700 Goat anti-mouse IgG antibody (1:5000 dilution) in the blocking buffer, for 1 hour on a rocking platform. Following another wash with 1X TBST buffer, repeated 5-6 times for 5 minutes each, the blot was imaged using a BioRad Gel Dock system to measure the expression levels of the target proteins relative to the housekeeping genes.

### TopFlash-FopFlash assay

10,000-15,000 cells per well were seeded in a white-bottom 96-well plate using 0.1 mL of DMEM supplemented with 10% FBS and 1% penicillin/streptomycin. The plate was then incubated at 37°C with 5% CO_2_ for 18-20 hours. For transfection, the TOPFlash plasmid (90 ng/well) or the FOPFlash plasmid (90 ng/well) as a negative control, along with the Renilla-luciferase plasmid (12 ng/well) as a normalization control were introduced to cells. Lipofectamine 2000 transfection reagent (0.5 µL/well) and serum-reduced Opti-MEM media were used for the transfection process. The lipofectamine and plasmid stocks were diluted in Opti-MEM to make a total volume of 20 µL of the plasmid-lipoplex solution that was added to each well (10 µL for each plasmid-lipoplex). The post-transfection total volume per well was approximately 120 µL. After transfection, cells were incubated for 6-8 hours. Subsequently, 80 µL of media was aspirated from each well and replaced with 50 µL of fresh DMEM media supplemented with 10% FBS and 1% penicillin/streptomycin. Next, the cells were treated with 10 µL of LNP formulations, appropriately diluted in 1X PBS to achieve different concentrations. The plate was incubated for an additional 24 hours. Following LNP treatment, media was aspirated, and the cells were washed twice with 1X PBS. Then, 20 µL of 1X passive lysis buffer (Promega Dual Glo kit) was added to each well and allowed to incubate for 2 minutes. For firefly luciferase measurement, the Firefly luciferase substrate (LAR II reagent, 100 µL/well, Promega Dual Glo kit, cat # E1960) was added to each well, and luminescence was measured using a microplate reader (TECAN Infinite M1000 Pro). Finally, the 1X Renilla-luciferase substrate mixed with a Stop & Glo buffer (100 µL/well, Promega Dual Glo kit, cat # E1960) was added, and the Renilla-Luminescence was measured to normalize the TOPFlash-Firefly luminescence.

### MTS cell viability assay

10,000 cells/well were seeded in a round-bottom transparent 96-well plate using 100 µL of DMEM/well supplemented with 10% FBS and 1% penicillin/streptomycin. The plate was incubated for 24 hours. Following this, 60 µL of media was removed from each well and 50 µL of fresh media was added along with 10 µL of LNP formulation, appropriately diluted to achieve the desired concentration. The plate was incubated again for different durations, specifically 24 hours, 48 hours, and 72 hours. After the specified incubation period, media was aspirated, and each well was washed twice with sterile 1X PBS. Next, 100 µL of dye-free Fluorbrite DMEM media containing 5% (5 µL) MTS reagent premixed in it was added to each well, followed by another 3 hour incubation. The absorbance was measured using a microplate reader (TECAN Infinite M1000 Pro) at a wavelength of 490 nm to assess cell viability and proliferation.

### In Vivo Biodistribution of LNP-encapsulated PepTACs

All animal experiments were conducted in accordance with an approved IACUC protocol (#2019-0063). Twenty-eight female SHK1 hairless mice (vendor -Charles River Laboratories, Elite strain 477) aged 8-11 weeks were housed in a clean facility with 12-hour light/dark cycles and fed alfalfa-free chow ad libitum.

Biodistribution studies were conducted using an adaptation of a previously published method.^70^ For biodistribution studies, mice (n=5 per group) were administered either 5 mg/kg free PepTAC labeled with 50% Cy5.5 or 5 mg/kg LNP-encapsulated PepTAC labeled with 50% Cy5.5 via tail vein injection (approximately 100 µL injection using a 26 G needle). 1X PBS (n=3, 100 µL injection) was used as a negative control. Mice were imaged using the IVIS Spectrum In Vivo Imaging Platform (Perkin Elmer, USA). To do this, mice were anesthetized using 3.5% isoflurane before transfer to the imaging stage of the IVIS Spectrum (Perkin Elmer, USA) maintained at 37 °C. During imaging, mice were maintained at 2% isoflurane anaesthesia. Mice were imaged in both supine and prone orientations at 1 and 24 hours. Images were obtained using a spectral unmixing sequence defined for Cy5.5. Following the final timepoint, mice were euthanized using 3.5% CO_2_ followed by cervical dislocation. The liver, spleen, kidneys, heart, lungs, brain, and GI tract were harvested and imaged ex vivo using the IVIS Spectrum (Perkin Elmer, USA).

To determine cellular-level biodistribution to the liver, 3 x 5 mm biopsy punches were obtained from separate lobes of the liver (excluding the caudate lobe). Samples were placed in histology cassettes and fixed in 10% (vol/vol) neutral buffered formalin for 48 hours prior to transfer to 1X PBS. Fixed samples were dehydrated and embedded in paraffin, and 4 µm sections were prepared for microscopy analysis. For confocal imaging, the slides were treated with Hoechst for nuclei-staining, and representative images from specific z-sections were captured using an oil-immersion 40x lens. In the captured images, blue channel emission shows the spatial distribution of the nuclei at different cells in the tissue sections while the red channel emission shows the Cy 5.5-labeled payloads.

To compare the organ distribution seen at 24 hours to that observed at 1 hour, female SKH1 mice (n=5 per group) were administered either 5 mg/kg free PepTAC labeled with 50% Cy5.5 or 5 mg/kg LNP-encapsulated PepTAC labeled with 50% Cy5.5 via tail vein injection (approximately 100 µL injection using a 26 G needle). After 1 hour, mice were sacrificed and harvested as described above. Ex vivo imaging of the organs was performed, and liver samples processed for histology. Samples were fixed in 10% (vol/vol) neutral buffered formalin for 48 hours followed by transfer to 1X PBS. Samples were prepared for microscopy as described above. To analyze the relative biodistribution of free PepTAC and LNP-encapsulated PepTAC to major organs, the Living Image Software (Perkin Elmer, USA) was used to draw custom regions of interest (ROIs) around images of organs taken ex vivo. The total radiant efficiency of each ROI was normalized to the pixel area of the ROI to obtain the average radiant efficiency.

### Assessment of in vivo protein degradation via LNP-encapsulated PepTACs

Thirty female BALB/c mice aged 6-8 weeks were housed in a clean facility with 12-hour light/dark cycles and fed ad libitum. Animals were identified via ear punch. To assess the activity of the PepTAC mice (n=5 per group) were administered LNP-encapsulated PepTACs (against β-catenin) at a dose of 5 mg/kg via tail vein injection (approximately 100 µL injection using a 26 G needle), one dose at every three days for 10 days (four doses) On the 11th day, mice were euthanized using 3.5% CO_2_ followed by cervical dislocation. The livers were immediately excised and rinsed with ice-cold sterile PBS buffer, and 3 x 5 mm biopsy punches were obtained from separate lobes of the livers (excluding the caudate lobe). Samples were placed in histology cassettes and fixed in 10% (vol/vol) neutral buffered formalin for 48 hours prior to transfer to 1X PBS. The remaining liver tissue was snap frozen in liquid nitrogen and stored at -80 °C. These liver samples were homogenized with RIPA lysis buffer premixed with 1X protease and phosphatase-inhibitor cocktail and 1x EDTA (to prevent non-specific protease, metalloprotease, or phosphatase-mediated digestion) using a benchtop homogenizer (VWR VDI 12 Homogenizer). Samples were centrifuged at 4000 rpm for 10 minutes at 4 °C, and the supernatants were passed through a 40 µm cell strainer to remove remaining tissue particulate. A standard BCA assay was performed to quantify the total protein concentration for each tissue homogenate. Western Blot was performed (loading: 20 µg total protein/well) with the tissue homogenates to assess the expression level of endogenous β-catenin as well as the housekeeping gene, β-tubulin.

## ACKNOWLEDGEMENTS

C.A.A acknowledges financial support from NSF CBET-1917285. C.R. was supported by an NSF GRFP fellowship (DGE-1650441). A.A. was supported by Fleming fellowship. J.A. was supported by the Ezra’s Bridge fellowship. The content is solely the responsibility of the authors and does not necessarily represent the official views of any of the above-mentioned funding agencies.

## AUTHOR CONTRIBUTIONS

S.G and C.A.A designed the research and wrote the manuscript. S.G., C.R., J.W., and H.W.C. performed the experiments. A.A. and J.A. generated research materials. C.A.A. supervised the research. All authors discussed the results and commented on the manuscript.

## ADDITIONAL INFORMATION

### Supplementary information

The online version contains supplementary material available at XXX.

**Peer review information**

**Reprints and permissions information**

