## Supplementary file for "Nanoparticle-mediated delivery of peptide-based degraders enables targeted protein degradation"

### Supplementary Figures

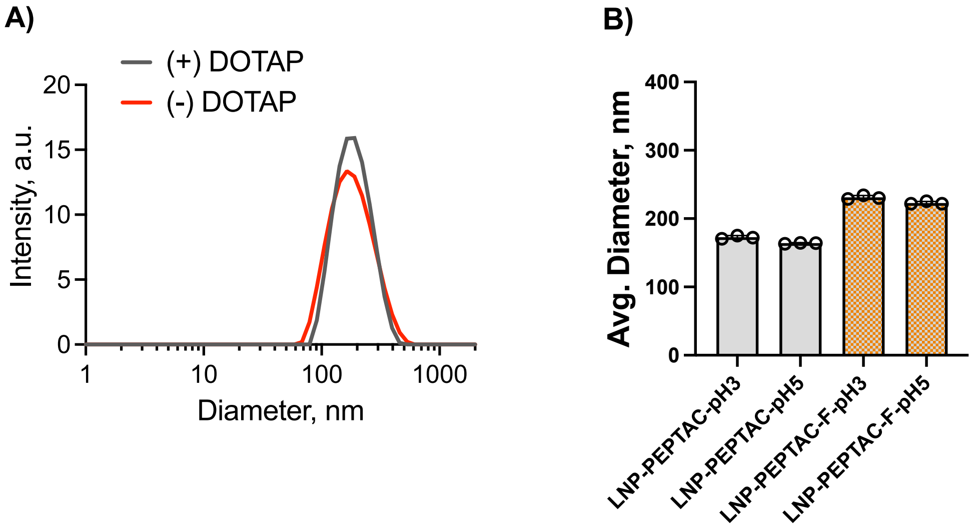

Figure S1| Dynamic light scattering

Dynamic light scattering of different iLNP:PepTAC formulations. A) LNP formulations (MC3:PepTAC 2w/w, pH formulation) with and without DOTAP. B) LNP formulations (MC3:PepTAC 2w/w, +DOTAP, ) at pH 3 and pH5, with and without a fluorophore on the PepTAC.

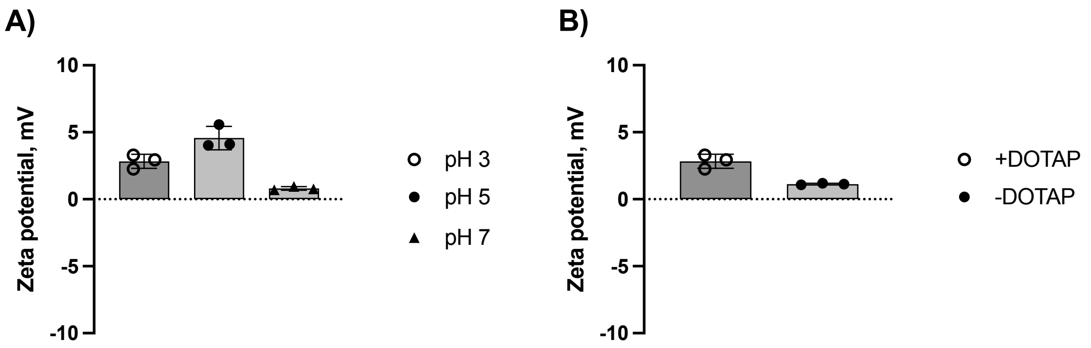

Figure S2| Zeta potential of iLNPs

Zeta potential measurements of iLNP:PepTAC formulations at A) different pH's, and B) with and without DOTAP.

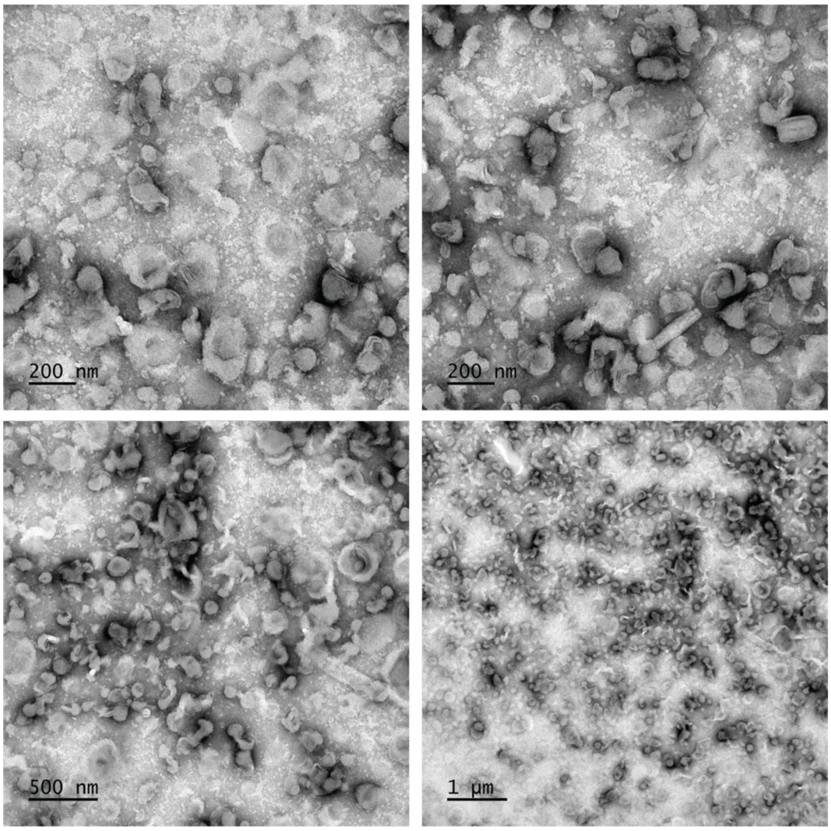

Figure S3| CyroTEM

CryoTEM of iLNP:PepTAC formulation (pH 3, MC3:PepTAC 2w/w with DOTAP)

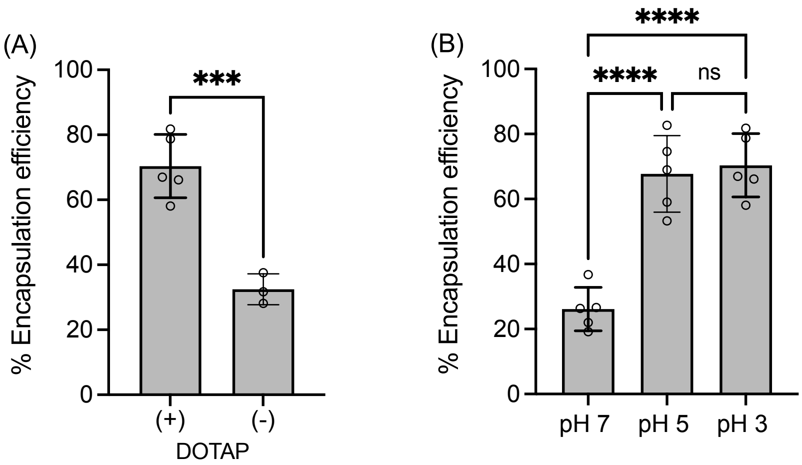

Figure S4| Encapsulation efficiencies of PepTAC-LNP formulations

Encapsulation efficiency of DOTAP LNP-PepTAC formulations (A) with and without DOTAP, and (B) with DOTAP at pH7, pH5, pH3

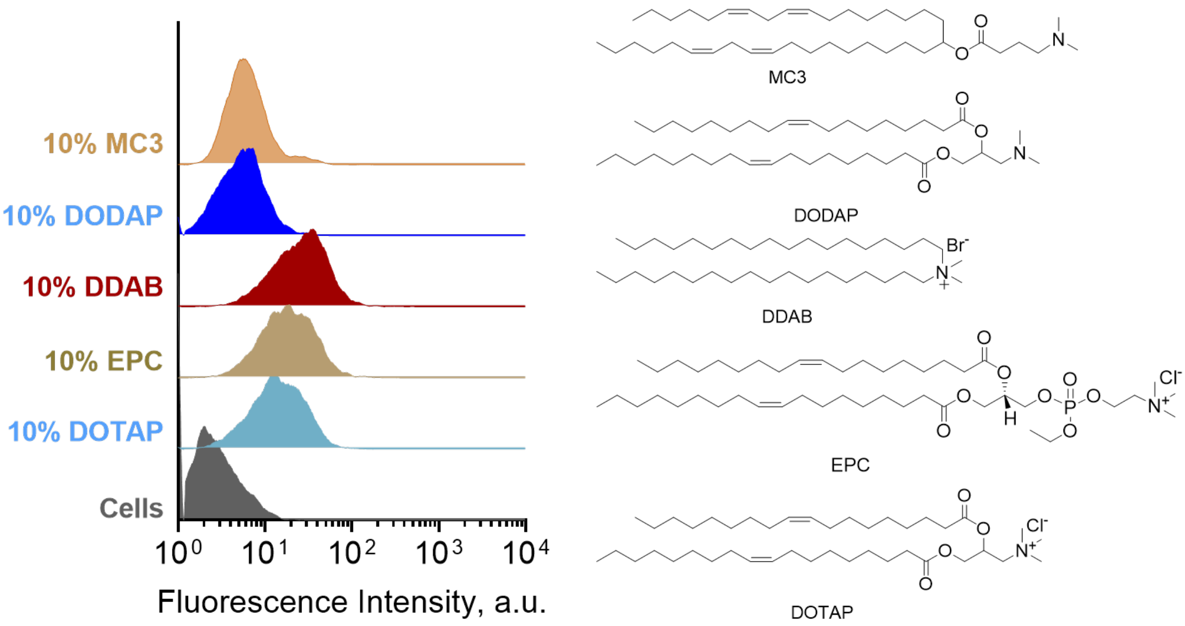

### Figure S5| Effect of structural analogs of DOTAP on cellular uptake

Effect of different structural analogs of DOTAP in the cellular uptake of fluorescein-labeled PepTAC in HeLa cells depicted in the histohrams obtained through flow cytometry. Careful observation of the subtle structural differences of different DOTAP-analogs shown in the right panel suggests that the permanently cationic nature of the 5th lipid is probably the most critical structural feature in order to obtain a good cellular uptake of labeled PepTAC.

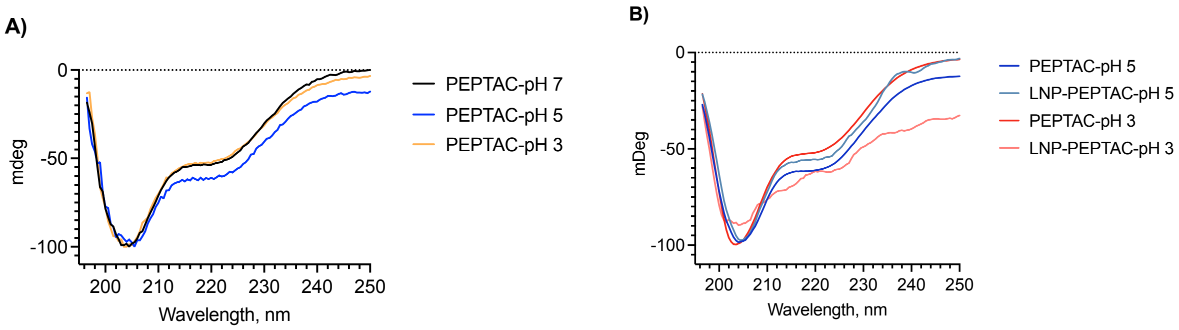

### Figure S6| Circular Dichroism

A) Circular Dichroism of PepTACs at different pH values used in the formulation. B) Circular Dichroism of LNP-PepTACs at different formulation pH values relative to PepTACs alone.

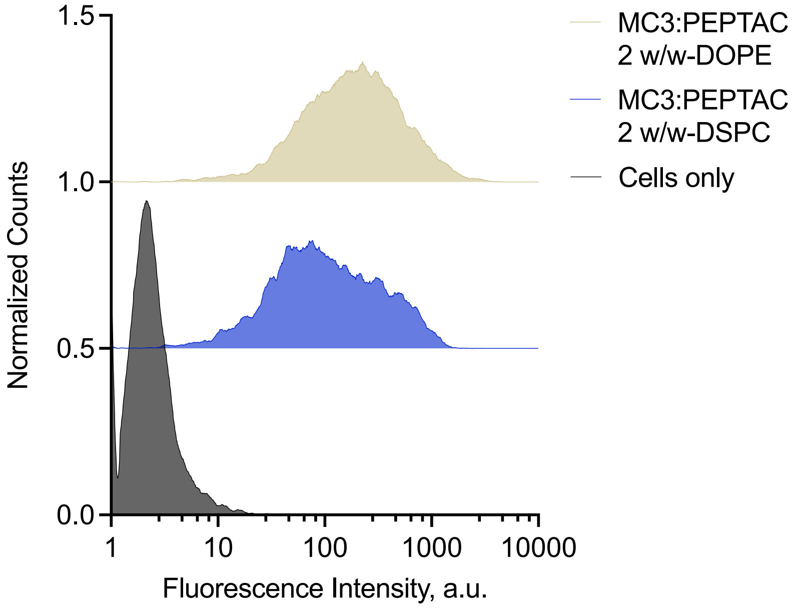

Figure S7| Effect of phospholipid on uptake in HeLa cells

Effect of DOPE vs DSPC in the LNP-PepTAC formulation. HeLa cells were transfected with the LNP-PepTAC formulation for 24hrs.

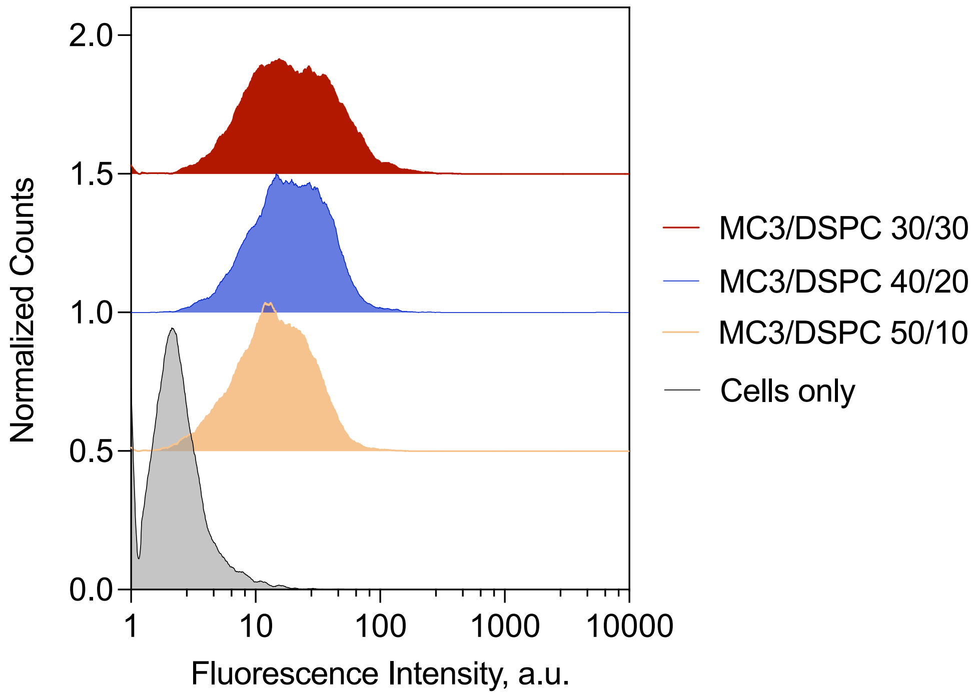

Figure S8| Effect of MC3: phospholipid on uptake in HeLa cells

Effect of MC3: DSPC ratio in the LNP-PepTAC formulation on uptake in HeLa cells

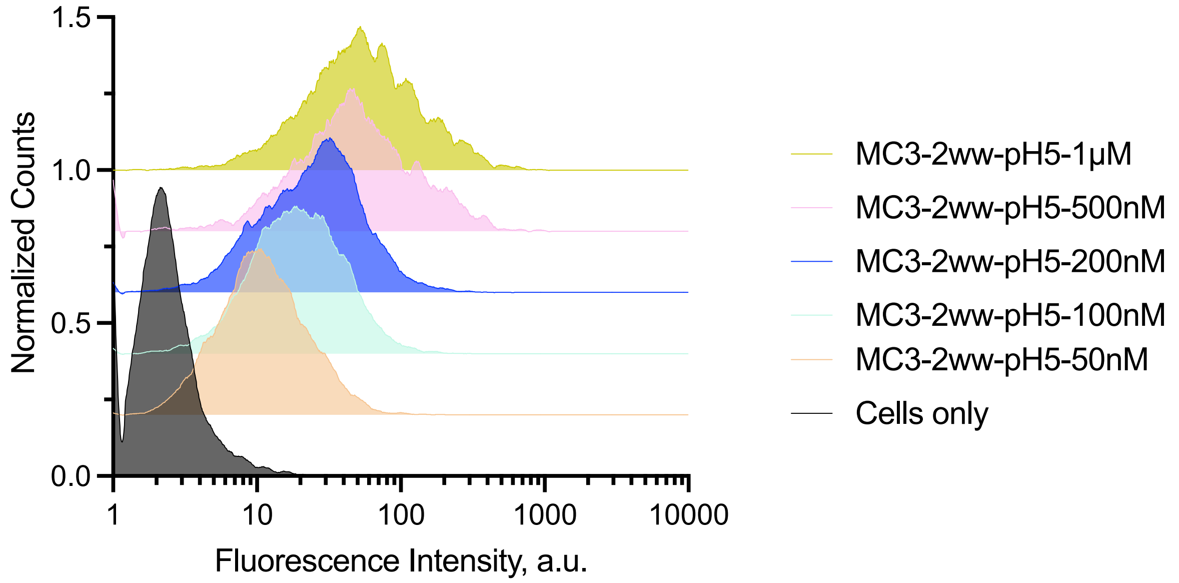

Figure S9| Dose dependent uptake at pH5 in HeLa cells
Dose dependent uptake of MC3 LNP-PepTACs formulated at pH5 in HeLa cells

A)

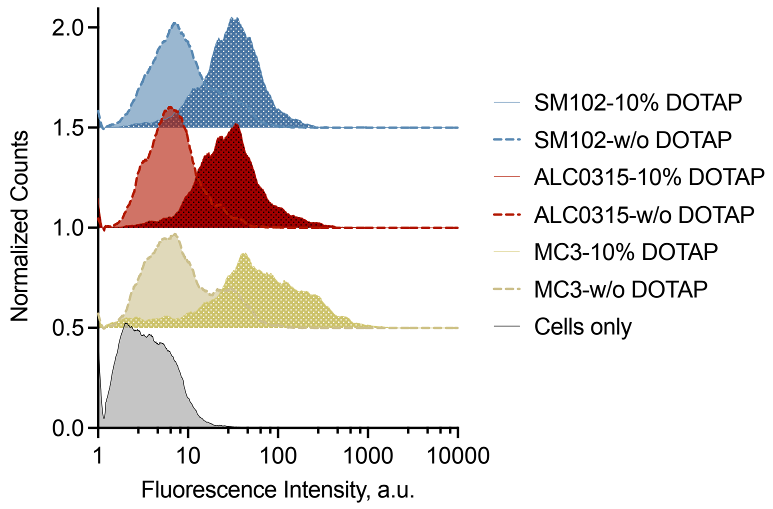

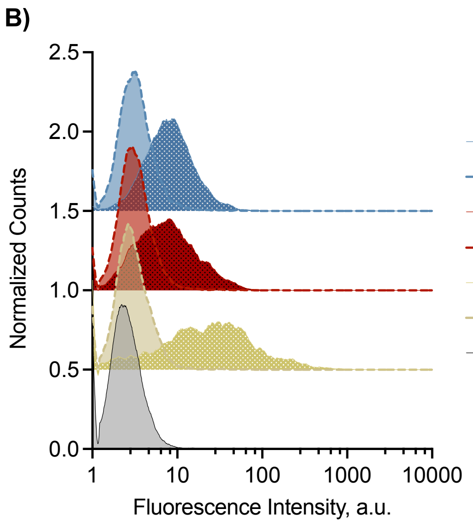

Figure S10| Effect of ionizable lipid composition and DOTAP

Effect of different ionizable lipids in A) SK-BR-3, and B) SKOV-3 cells

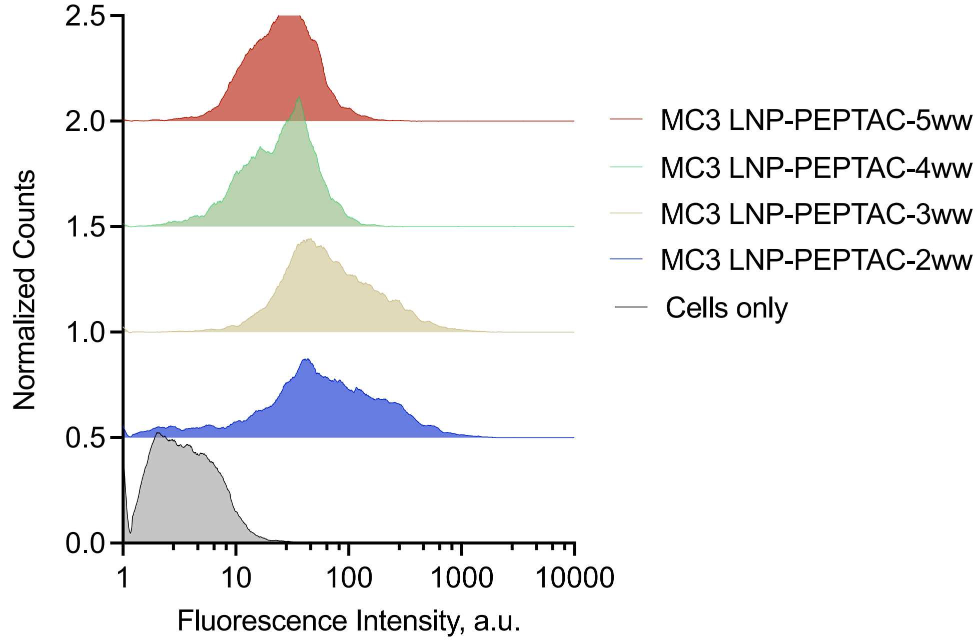

Figure S11| Effect of MC3:PepTAC wt/wt ratio on uptake in SK-BR-3 cells

Effect of MC3:PepTAC wt/wt ratio on SK-BR-3 cellular uptake

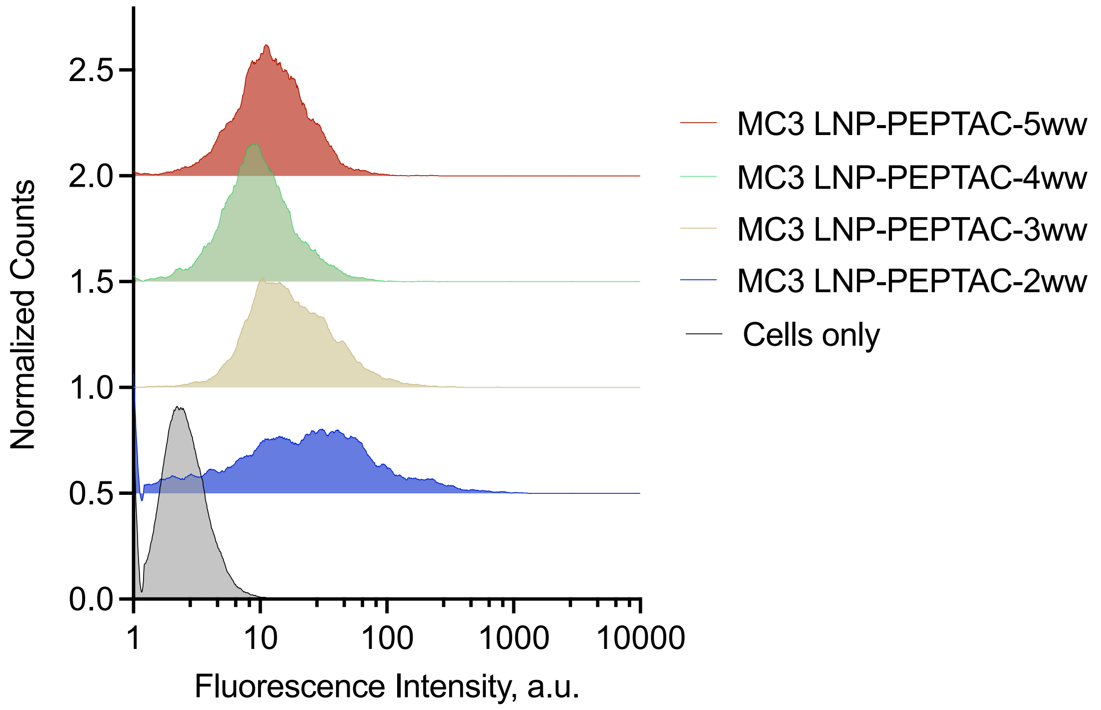

Figure S12| Effect of MC3:PepTAC wt/wt ratio on uptake in SKOV-3 cells

Effect of MC3:PepTAC wt/wt ratio on SKOV-3 cellular uptake

**A)**

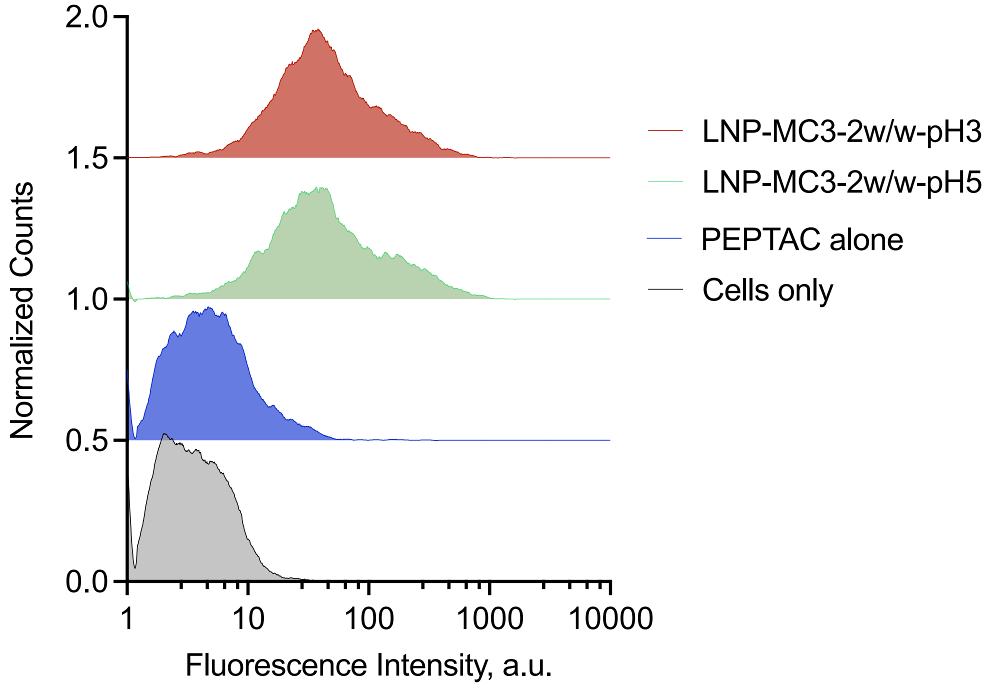

B)

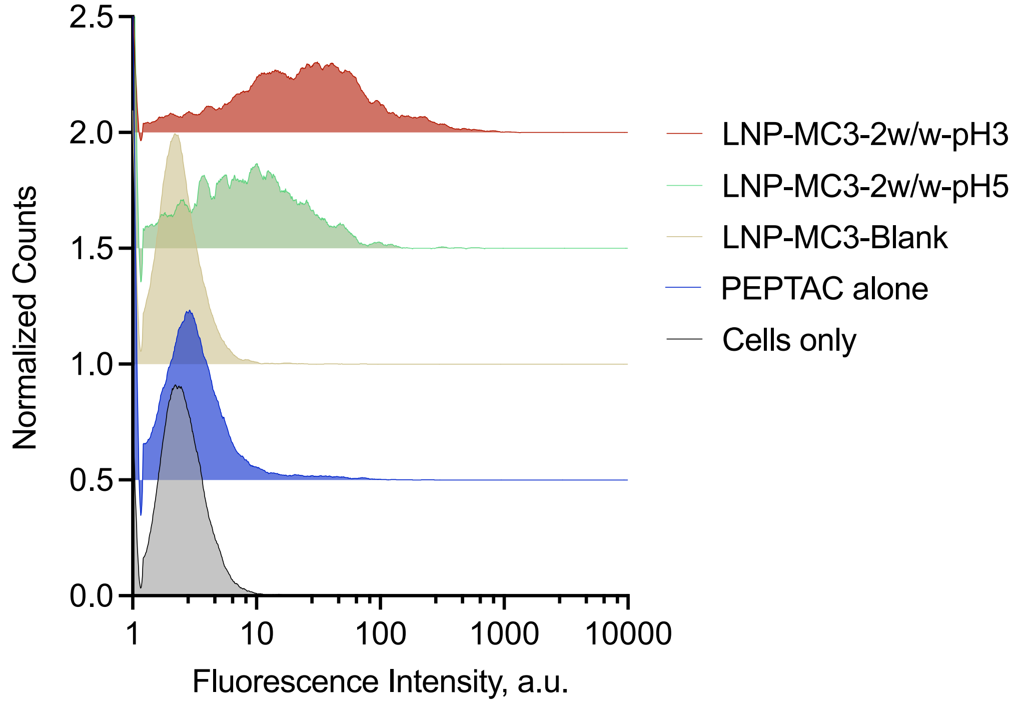

Figure S13| Effect of formulation pH on uptake in different cells
Effect of pH on uptake in SKBR-3 and SKOV-3 cells

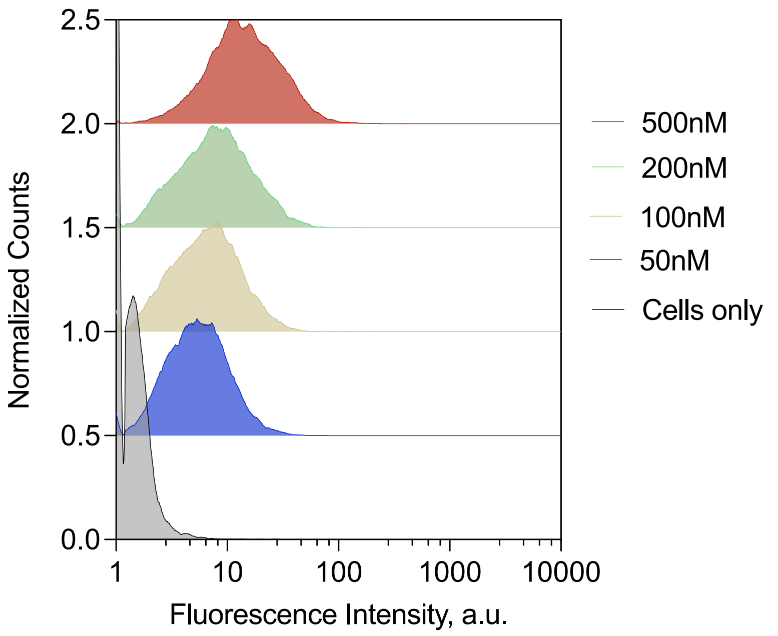

Figure S14| Dose-dependent uptake in DLD1 cells
Systematic dose-dependent cellular uptake of fluorescein-labeled $^{CR}$PepTAC in DLD1 cells

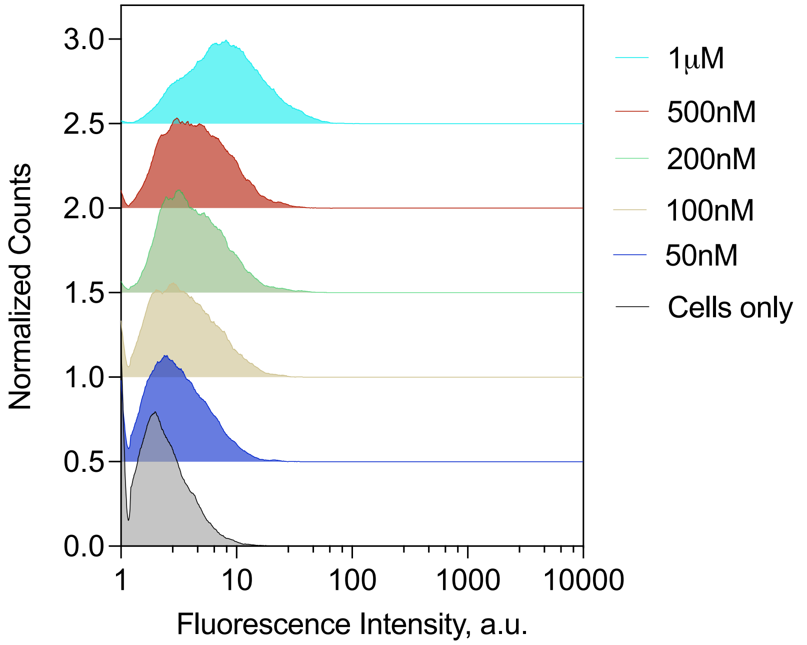

Figure S15| Dose-dependent uptake in HepG2 cells
Systematic dose-dependent cellular uptake of fluorescein-labeled $^{CR}$PepTAC in HepG2 cells

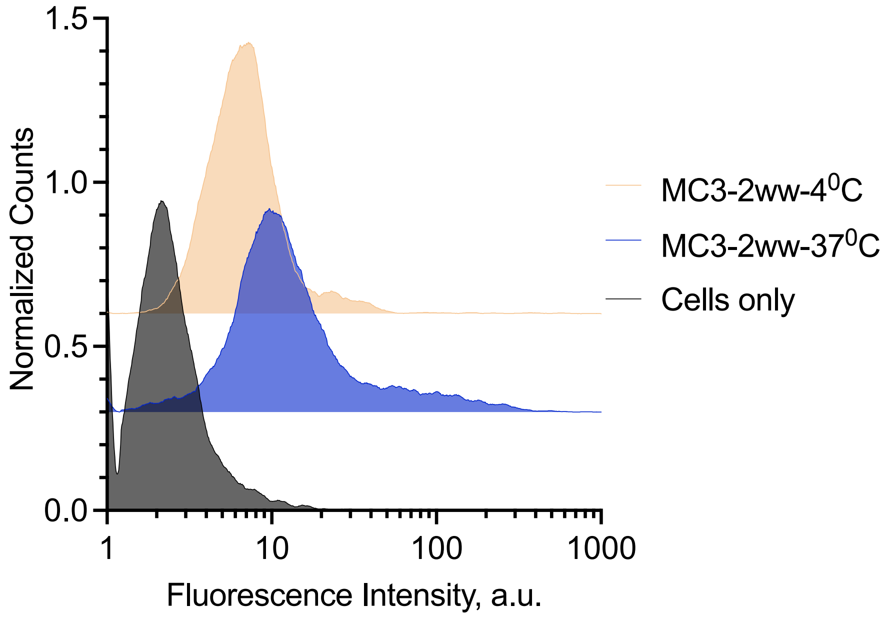

Figure S16| Effect of temperature on uptake
Temperature dependent uptake of LNP-PepTACs in HeLa cells (MC3:PepTAC 2wt/wt, 2h incubation with 500 nM PepTAC)

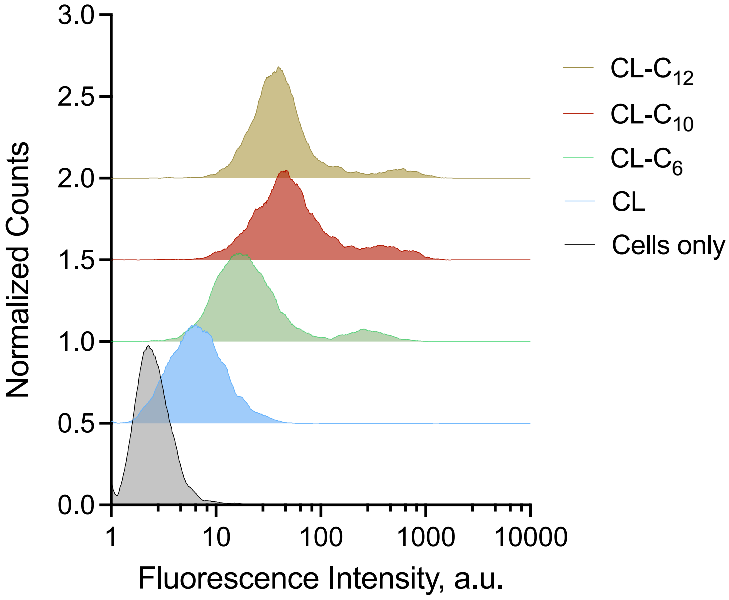

Figure S17| Cellular uptake of lipid modified CREPT ligand (CL) peptides in HeLa cells
Cellular uptake of lipid modified CL peptides of different lipid tail lengths (C6, C10 and C12) formulated with LNPs and transfected into HeLa cells.

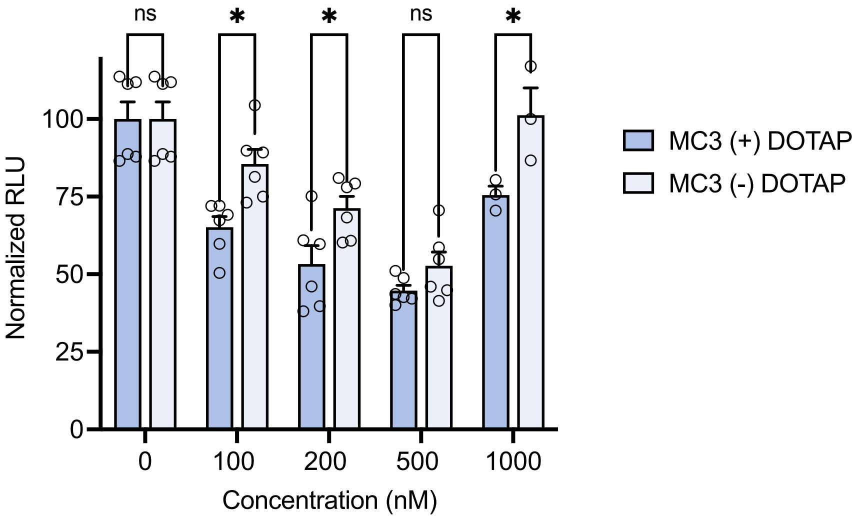

Figure S18| Functional effect of DOTAP in HeLa cells
Effect of DOTAP as a 5$^{th}$ lipid in the LNP formulation of $^{CR}$PepTAC. CREPT degradation was measured in HeLa cells transiently expressing Firefly luciferase-fused CREPT protein.

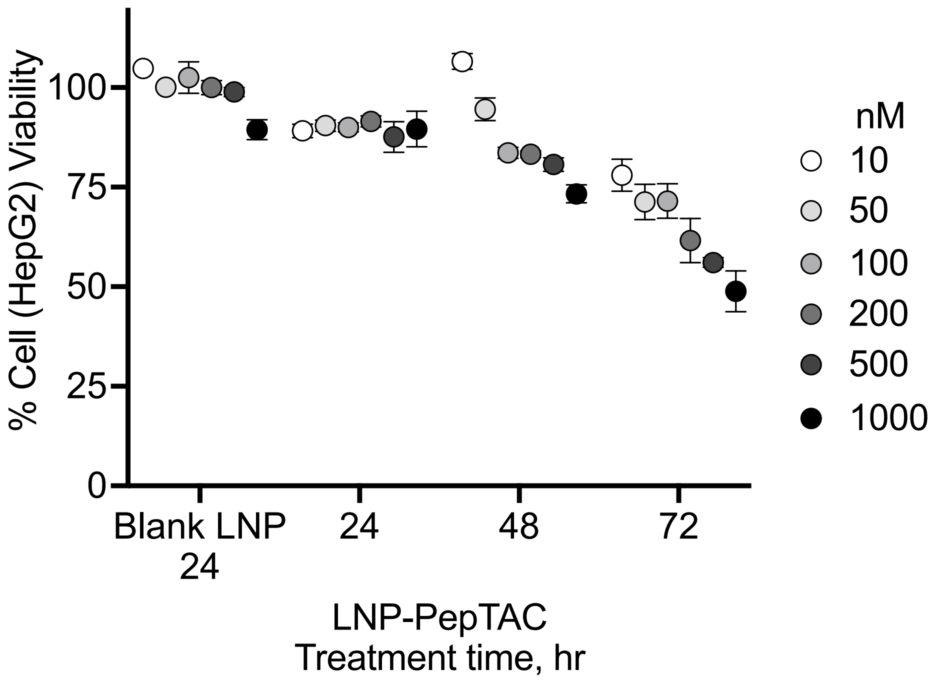

Figure S19| Effect of $^{CR}$PepTAC-LNP on HepG2 cell viability at different timepoints
Effect of LNP- $^{CR}$PepTAC treatments at different concentrations on Wnt-active HepG2 cell viability at 24, 48 and 72h. Cell viability was assessed via an MTS assay.

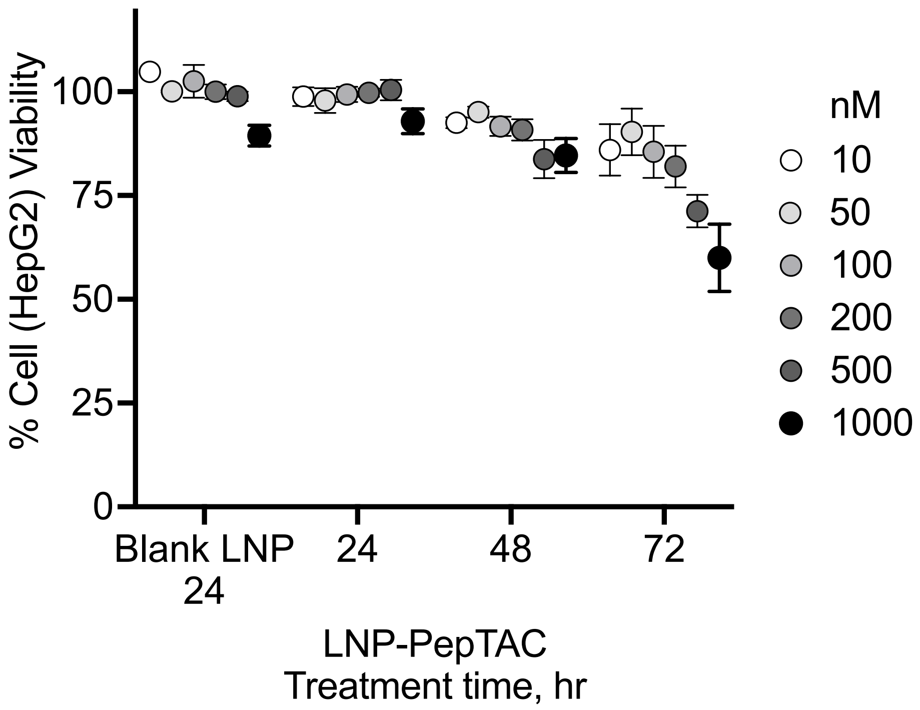

Figure S20| Effect of $^{\beta Cat}$PepTAC-LNP on HepG2 cell viability at different timepoints
Effect of LNP- $^{\beta Cat}$PepTAC treatments at different concentrations on Wnt-active HepG2 cell viability at 24, 48 and 72h. Cell viability was assessed via an MTS assay.

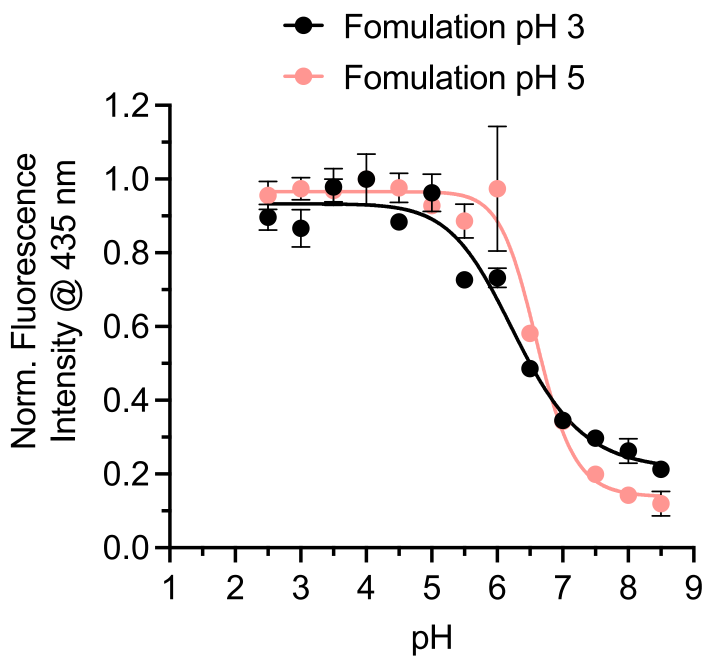

Figure S21| Apparent pKa of $^{CR}$PepTAC-LNP (MC3)

Apparent pKa (using the TNS assay) of MC3 PepTAC-LNPs formulated at pH3 and pH5 (MC3: PepTAC 2w/w).

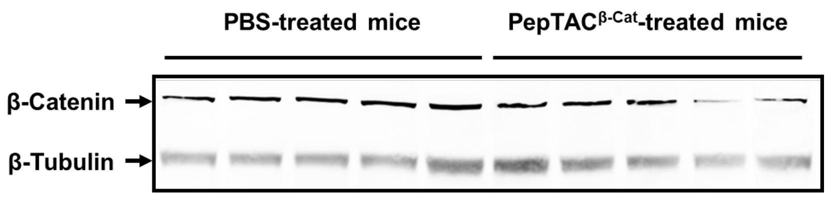

Figure S22| Western Blot image showing β-Catenin degradation in mice liver homogenates

Representative Western Blot image showing β-Catenin expression level in mice treated with saline versus mice treated with $^{βCat}$PepTAC encapsulation inside LNP. n = 5 per group.

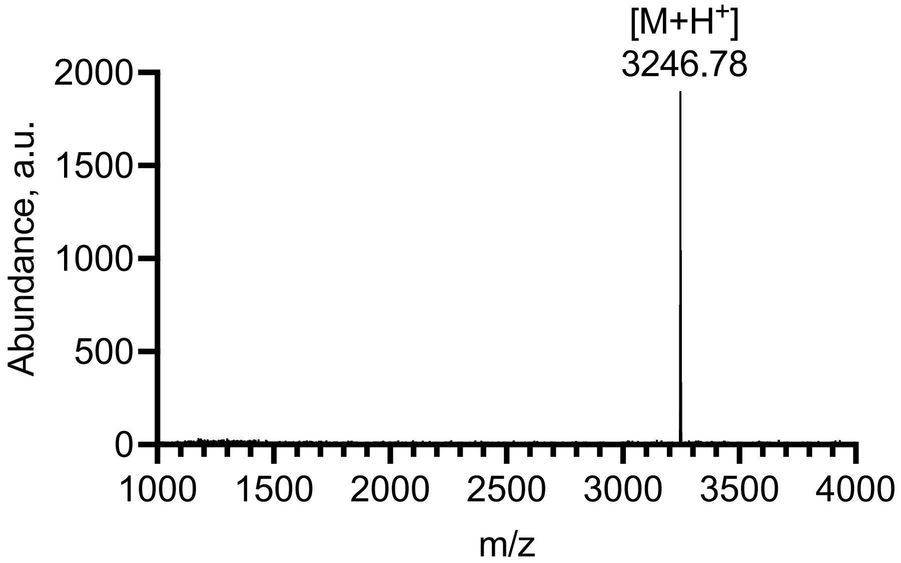

Figure S23| $^{CR}$PepTAC characterization
MALDI MS characterization of $^{CR}$PepTAC.

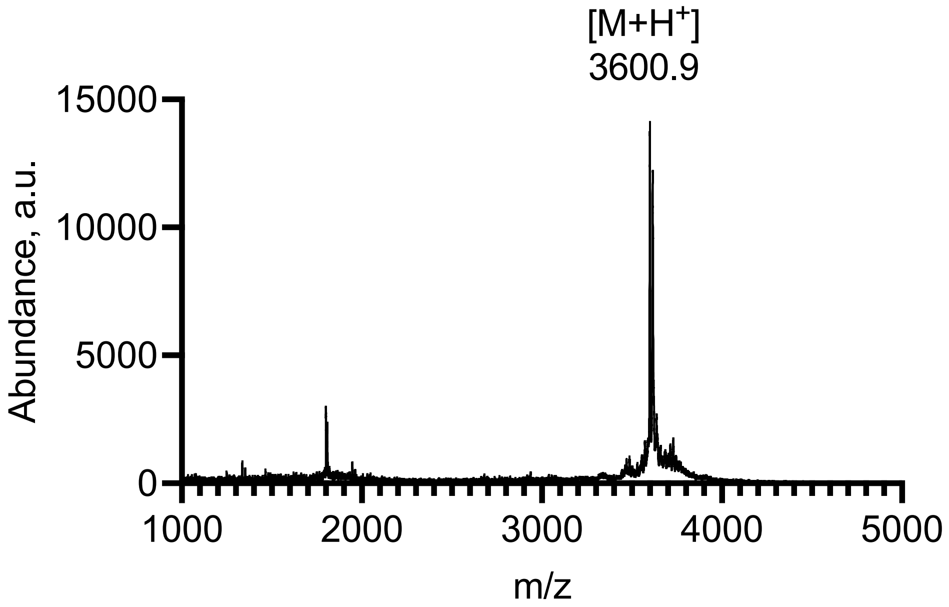

Figure S24| $^{\beta Cat}$PepTAC characterization
MALDI MS characterization of $^{\beta Cat}$PepTAC.

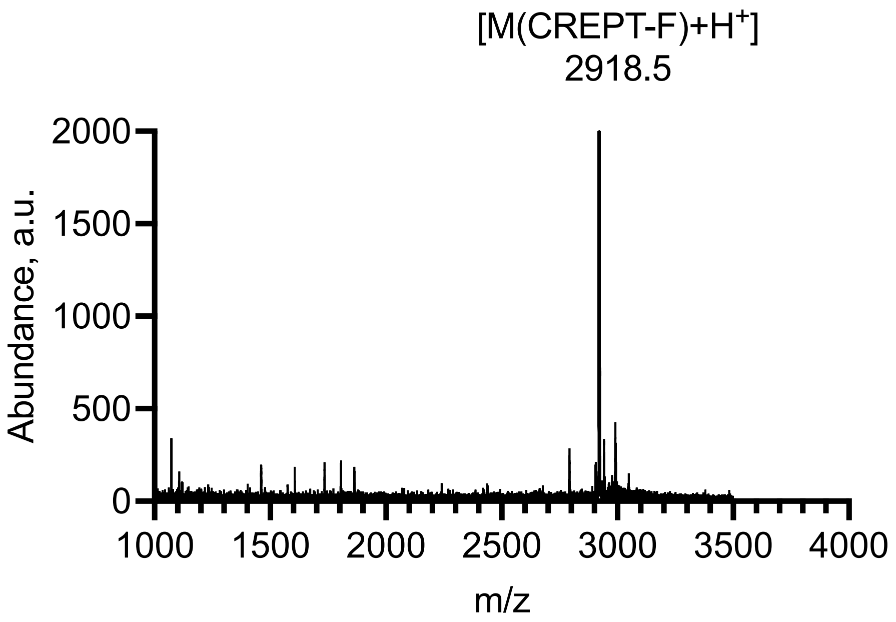

Figure S25| Fluorescein labeled CREPT Ligand
MALDI MS characterization of fluorescein labeled CREPT ligand (label at N-terminal)

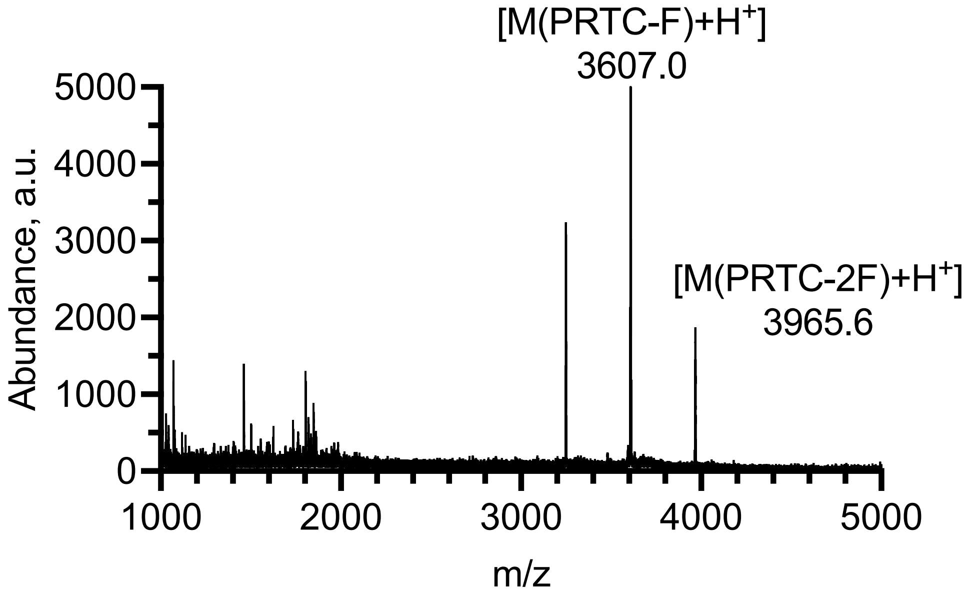

Figure S26| Fluorescein labeled $^{CR}$PepTAC
MALDI MS characterization of fluorescein labeled $^{CR}$PepTAC

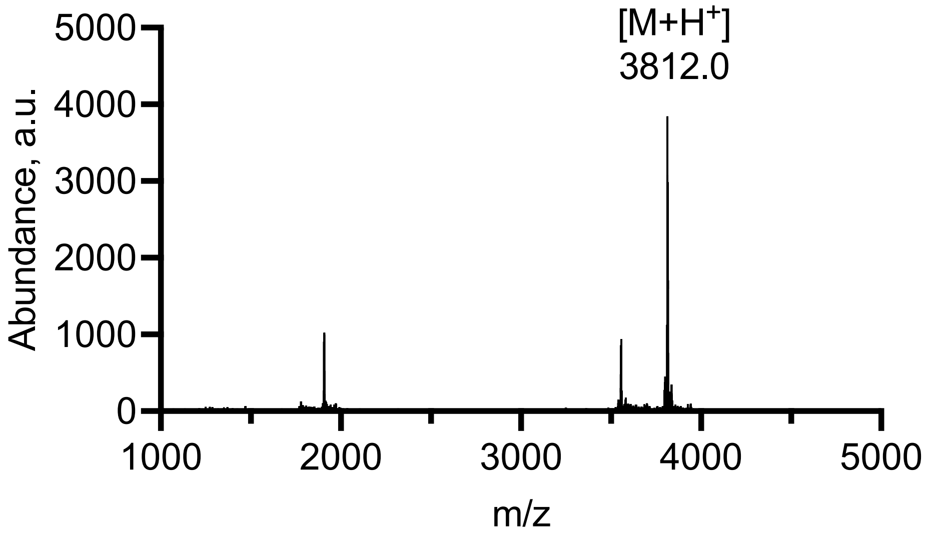

Figure S27| Cy5.5 labeled $^{CR}$PepTAC
MALDI MS characterization of Cy5.5. labeled $^{CR}$PepTAC

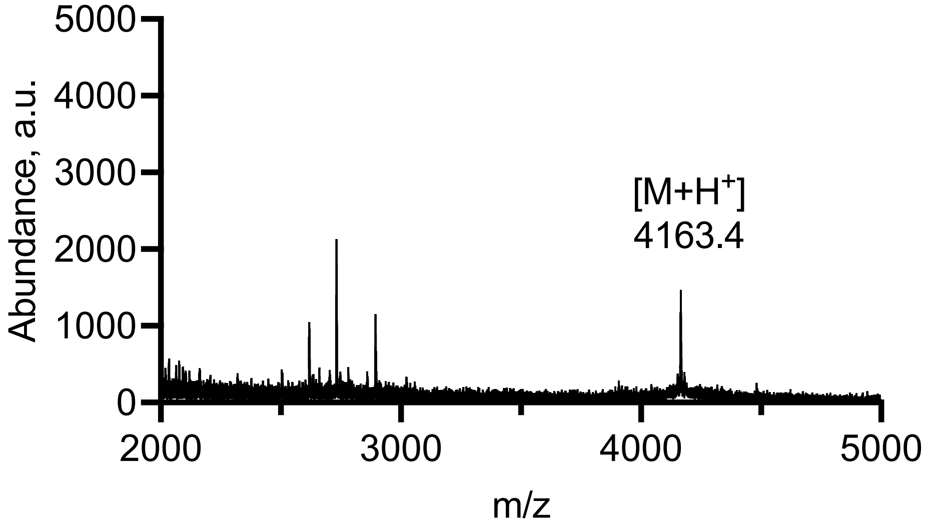

Figure S28| Cy5.5 labeled $^{\beta Cat}$PepTAC
MALDI data for the characterization of Cy5.5. labeled $^{\beta Cat}$PepTAC.

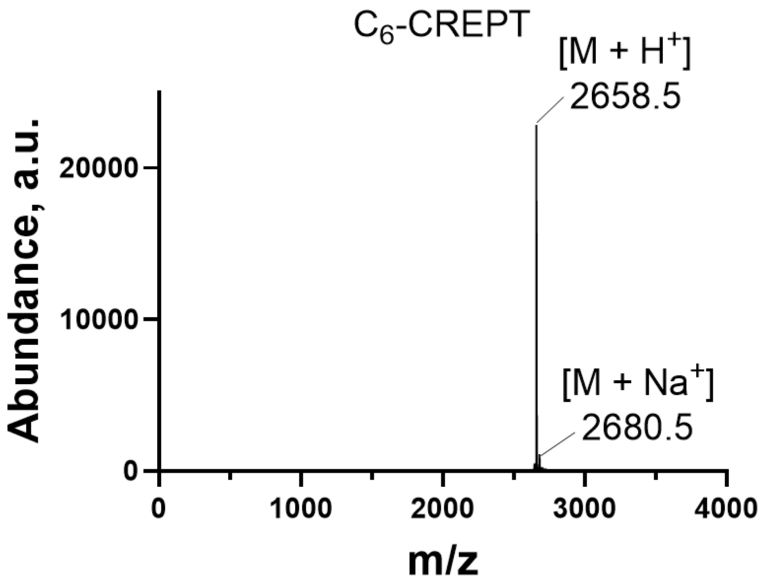

Figure S29| C6-CREPT lipidated peptide construct
MALDI data for the characterization of C6-CREPT lipidated peptide construct

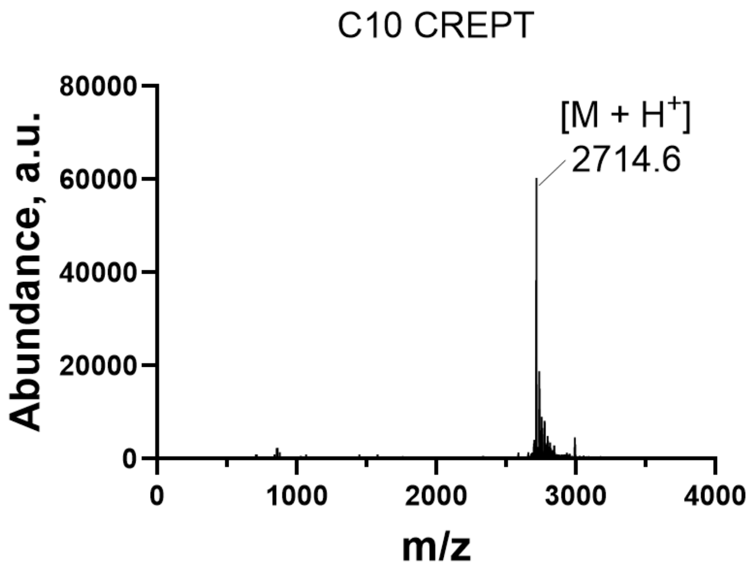

Figure S30| C10-CREPT lipidated peptide construct
MALDI data for the characterization of C10-CREPT lipidated peptide construct

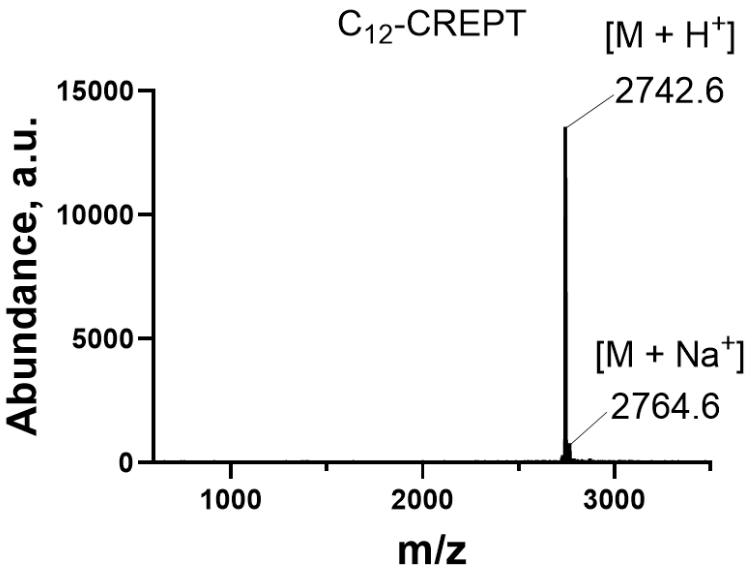

Figure S31| C12-CREPT lipidated peptide construct
MALDI data for the characterization of C12-CREPT lipidated peptide construct

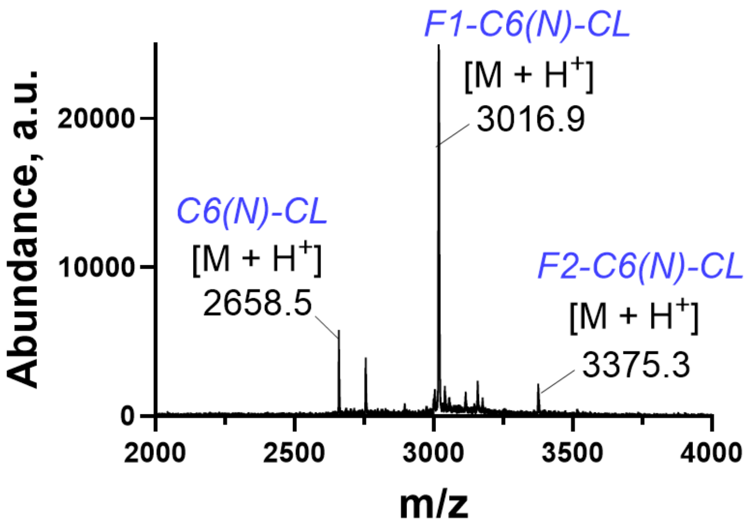

Figure S32| Fluorescein-conjugated C6(N)-CL lipidated peptide (CREPT ligand) construct
MALDI data for the characterization of Fluorescein-conjugated C6(N)-CL lipidated peptide (CREPT ligand) construct showing the formation of both singly and doubly fluorescein-conjugated lipidated peptide (CL) construct

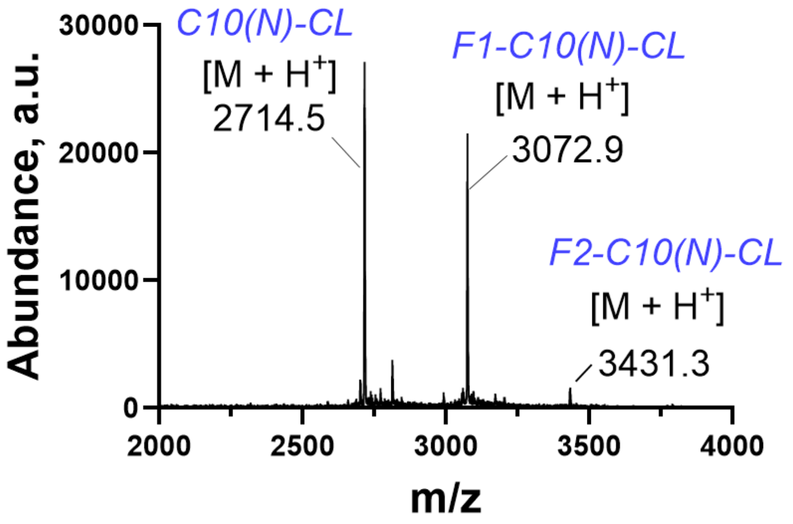

Figure S33| Fluorescein-conjugated C10(N)-CL lipidated peptide (CREPT ligand) construct
MALDI data for the characterization of Fluorescein-conjugated C10(N)-CL lipidated peptide (CREPT ligand) construct showing the formation of both singly and doubly fluorescein-conjugated lipidated peptide (CL) construct

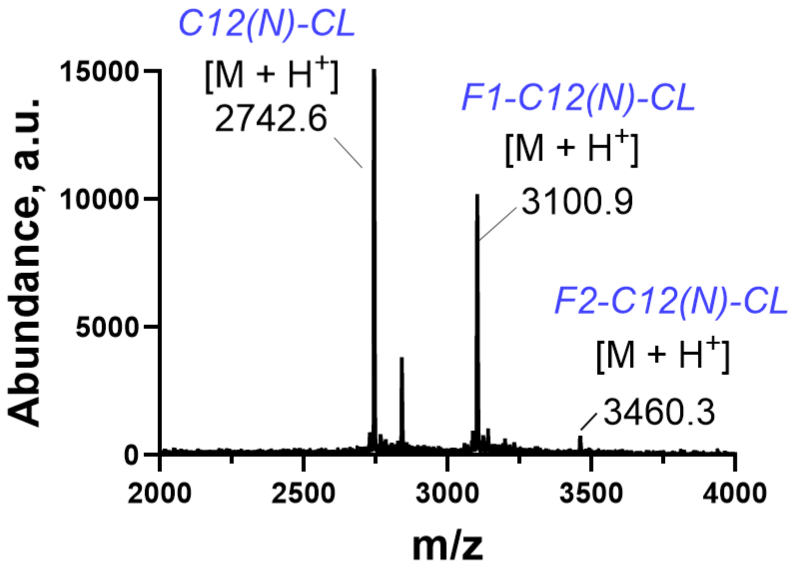

Figure S34| Fluorescein-conjugated C12(N)-CL lipidated peptide (CREPT ligand) construct
MALDI data for the characterization of Fluorescein-conjugated C12(N)-CL lipidated peptide (CREPT ligand) construct showing the formation of both singly and doubly fluorescein-conjugated lipidated peptide (CL) construct

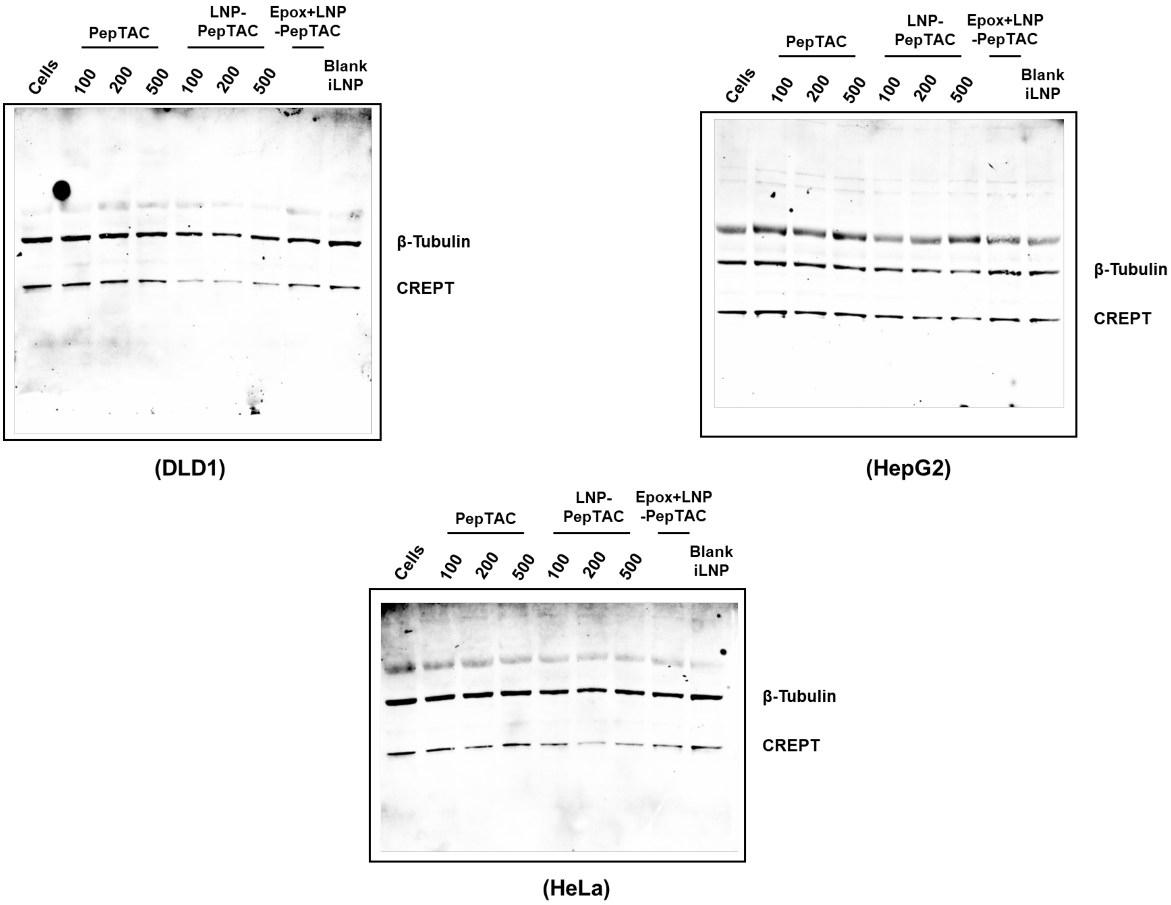

Figure S35| Western Blot raw data for CREPT degradation in three different cell lines

Raw Western Blot images showing CREPT degradation in DLD1, HepG2 and HeLa cells following treatment with $^{CR}$PepTAC-LNP.

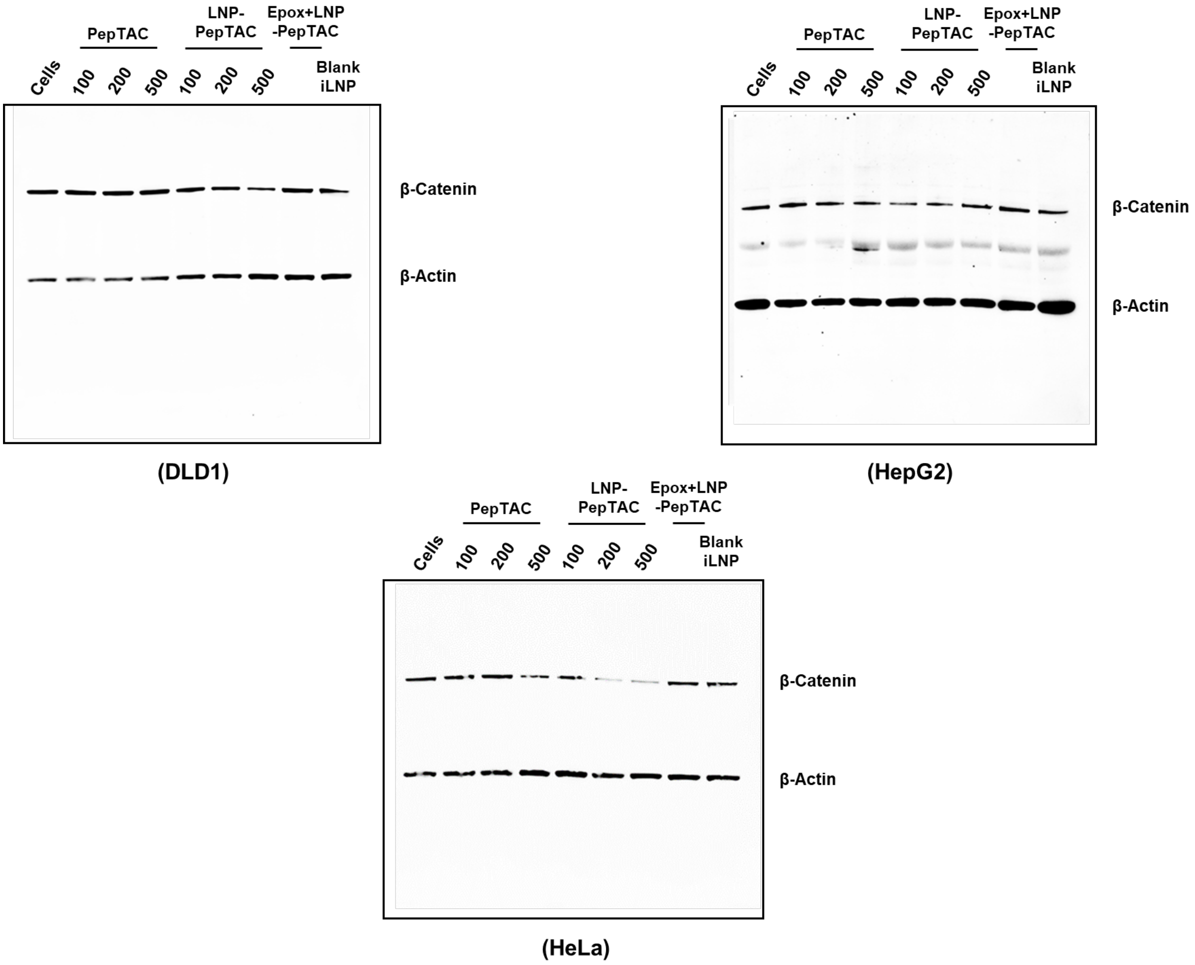

Figure S36| Western Blot raw data for β-Catenin degradation in three different cell lines
Raw Western Blot images showing β-Catenin degradation in DLD1, HepG2 and HeLa cells following treatment with $^{\beta Cat}$PepTAC-LNP.

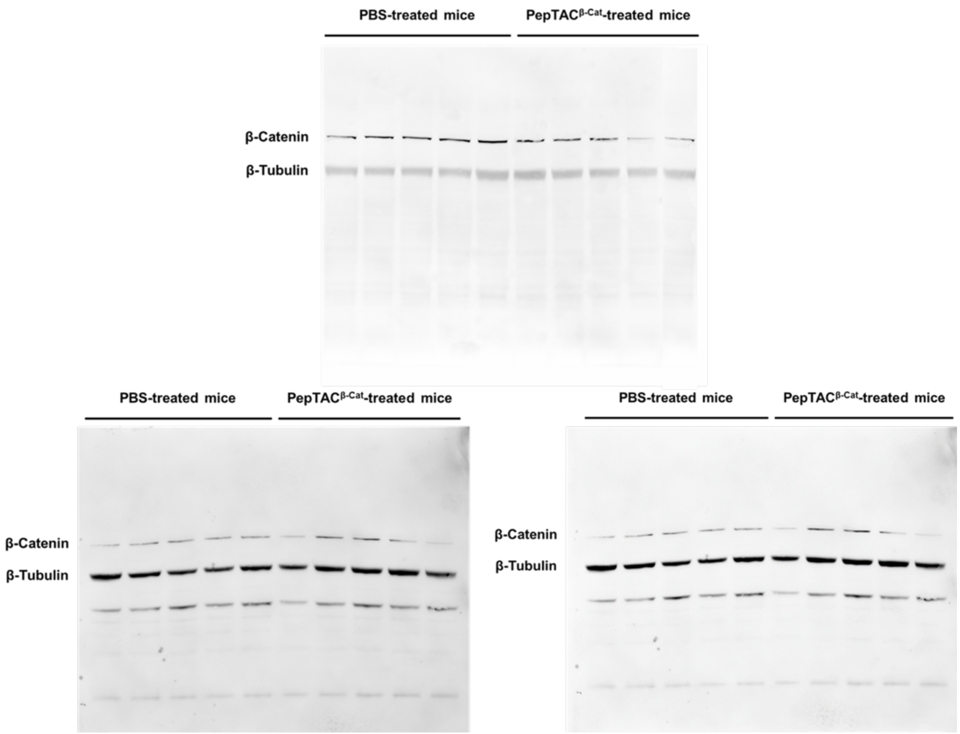

Figure S37| Western Blot raw data for β-Catenin degradation in mice liver homogenates

Western Blot images (three replicates) showing β-Catenin expression level in mice treated with saline versus mice treated with $^{\beta Cat}$PepTAC encapsulation inside LNP. n = 5 per group.
